# Mice sense Moon and Sun

**DOI:** 10.64898/2026.08.17.745150

**Authors:** Warsha Barde, Alexander Grayver, Annette E. Rünker, Mariko Izumo, Victoria A Acosta-Rodríguez, Joseph S Takahashi, Gerd Kempermann

## Abstract

Life on earth has always been exposed to the fluctuating Earth’s magnetic field, but a magnetic sense affecting behavior has been debated for mammals. We here report that mice, kept under constant laboratory conditions, showed fluctuations in spontaneous behavioral activity with a periodicity of ∼14 and ∼28 days. This was confirmed in nine cohorts from four facilities on two continents, covering 3 to 41 months. Such oscillations were also maintained in brain *Bmal1* knockout mice lacking circadian rhythms, suggesting independence of the circadian clock. The behavioral activity peaked around full and new Moon, and showed a strong alignment with the periodic geomagnetic fluctuations originating in the Earth’s iono- and magnetosphere that are modulated by solar rotation and the orbital motion of the Moon. In the ultradian range, this alignment persisted in CRY1/2 knockout mice, suggesting that solar–lunar-driven geomagnetic fluctuations can modulate behavior rhythms independently of CRY1/2.

## Introduction

Biological rhythms enable organisms to anticipate and adapt to predictable environmental changes. Among these, circadian rhythms with a ∼24-hour period are the most extensively characterized, driven by a cell-autonomous and self-sustaining transcription-translation feedback loop of clock genes. In mammals, the central circadian clock located within the suprachiasmatic nucleus (SCN) that regulates rhythmic behavior in sleep/wake and feeding/fasting cycles is primarily entrained by the Earth’s solar day (24 h) ^1,2^.

Beyond the circadian timescale, organisms express rhythms at both higher and lower frequencies. A range of biological rhythms with frequencies higher than the circadian cycle are collectively termed as ultradian rhythms. These include well-documented patterns such as ultradian sleep cycles ^3,4^, feeding bouts ^5,6^, locomotor activity ^7^, and pulsatile hormone release ^8,9^. While some are tightly linked to known physiological or behavioral processes, many ultradian oscillations remain poorly understood in terms of their underlying mechanisms ^10,11^. At the other end of the spectrum are slower biorhythms, with periods exceeding 24 hours, broadly categorized as infradian rhythms. These include lunar, seasonal, annual, and biannual cycles each reflecting adaptations to predictable, long-period environmental changes. The expression of seasonal and annual rhythms is well characterized and is primarily entrained by changes in photoperiod and temperature, which govern migration ^12,13^, hibernation ^14,15^, and reproduction ^12,16^ While tidal and lunar synchronization of behavior is well-established in marine and amphibious species ^17,18^, recent evidence suggests that even in humans, menstrual cycles can transiently align with both the luminance and orbital cycles of the Moon ^19^.

In laboratory rodents, the presence of infradian rhythms remains remarkably underexplored with few exceptions ^20–23^. This gap stems partly from a prevailing focus on circadian mechanisms, but also from methodological limitations: traditional behavioral paradigms are short in duration, limited in resolution, and often insensitive to long-period oscillations. Furthermore, laboratory animals are typically housed under constant lighting, feeding, and temperature regimes, thought to be insulating them from natural long-period zeitgebers.

With the purpose of investigating gene – environment interactions in shaping individual behavior and brain plasticity among genetically identical animals in a shared enriched environment, we developed a longitudinal framework termed the “Individuality Paradigm” ^24^. This approach uses the ColonyRack system - a custom-built, enriched housing environment - where group-housed mice can freely explore a network of 70 interconnected cages via RFID-tracked connector tubes. With a total floor area of 2.74 m² and spatial resolution at the level of individual cages, this setup enables continuous, high-resolution tracking of spontaneous behavior of individual mice across weeks to months.

## Results

### Long-term behavioral tracking reveals infradian rhythms

We performed an extensive meta-analysis of long-term (3 – 7 months) ColonyRack datasets from group-housed isogenic mice, which allow for continuous, non-invasive monitoring of socially housed mice over long time periods. Six datasets (D1–D6), recorded between 2021 and 2023, each comprising 33-40 mice and spanning 90-200 days, were included in this analysis. Behavioral dynamics were assessed using Roaming Entropy (RE), a quantitative metric reflecting the entropy of spatial location transitions that effectively captures patterns of exploratory behavior ^25,26^. RE ranges from 0 to 1, with lower values indicating that mice restricted their activity to only a few cages, and higher values indicating a more even distribution of activity and time across cages, reflecting broader spatial exploration. For each dataset, RE was computed at an hourly resolution and visualized as heatmap actograms averaged across animals (Fig. 1a). As expected, these actograms show clear circadian rhythmicity, showing activity synchronization to lights-on (6 AM) and lights-off (6 PM). Spectral analysis at hourly resolution across all datasets revealed dominant peaks at circadian (24 h) and ultradian (12 h, 6 h) periods, consistent with known high-frequency behavioral oscillations (Fig. 1c).

**Fig. 1.**
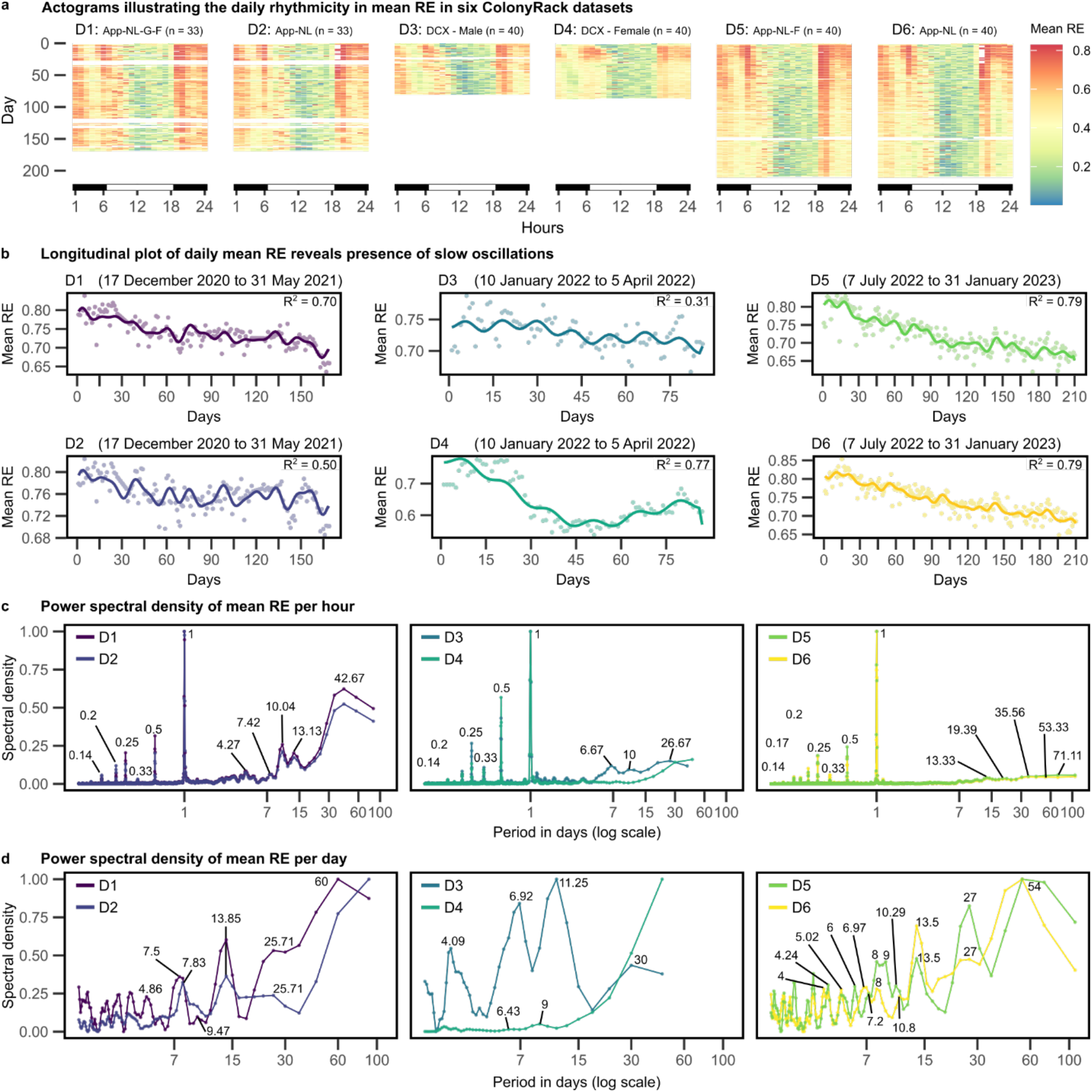
Longitudinal and spectral analysis of mean roaming entropy (RE) across six ColonyRack datasets reveals the presence of slow infradian rhythmicity in addition to ultradian and circadian rhythms. **(a)** Actograms with color-coded activations illustrating the daily rhythmicity in RE synchronized to lights-on (6 AM) and lights-off (6 PM) for six ColonyRack datasets D1-D6. Missing values are colored as white. **(b)** Longitudinal trends in daily mean RE suggest the presence of slower, infradian oscillations. The continuous bold line for each dataset represents a reconstructed waveform, generated by summing sine waves based on the dominant frequencies, amplitudes, and phases identified in the Fourier spectrum. Datasets were collected in three recording phases: D1–D2 over 166 days (17 December 2020 to 31 May 2021), D3–D4 over 86 days (10 January 2022 to 5 April 2022), and D5–D6 over 208 days (7 July 2022 to 31 January 2023). **(c)** Spectral analysis (normalized from 0 to 1) of mean hourly RE revealed circadian, ultradian, and infradian components in spontaneous activity. Numbers in the plot mark the peaks in days. **(d)** Spectral density plots of daily mean RE from D1-D6 show clustering of power at periodicities near multiples of 15 days, indicating a potential semi-lunar modulation. Numbers in the plot mark the peaks in days.

Visual inspection of time series of six datasets aggregated into daily means, however, also showed distinct slow, large-amplitude fluctuations in mean RE levels characterized by alternating periods of elevated and decreased values (Fig. 1b). This oscillatory pattern could be consistently observed across all datasets. Moreover, datasets recorded in tandem (D1–D2, D3–D4, D5–D6) revealed a striking synchronization of peaks and troughs, highlighting a remarkable temporal coordination in the occurrence of these oscillatory fluctuations. While the amplitudes of these peaks and troughs exhibited minor variations, the synchronized alignment hinted at clustering of activity at specific frequencies and possibly an underlying regulatory mechanism that orchestrates these rhythms, potentially stemming from shared environmental or physiological influences.

To quantify these slower rhythms, we conducted spectral analysis at daily resolution, which revealed consistent clustering of power at ∼15-day and ∼30-day periods across datasets (Fig. 1d). Additional spectral power was observed at multiples of ∼15 days, hinting at potential harmonic structure. To examine how frequency content varied over time, we used spectrograms, which confirmed clustering of power around ∼15 and ∼30 days (Fig. S1 a), supporting the presence of quasi-stable harmonic components at infradian scales.

These dynamics could not be explained by known environmental variables. All cohorts were maintained under stable 12:12 light-dark cycles, with controlled temperature, humidity, and *ad libitum* feeding. Weekly cage changes caused only a brief, transient increase in RE that resolved within a few hours (Fig. S2 a), and no alignment was found between the peaks or troughs of the long-period behavioral oscillation and cage-cleaning days or seasonal changes (Fig. S2 b).

### Behavioral alignment with lunar cycles

Given the persistence, periodicity, and coordination of these slow fluctuations, and their clustering near ∼15- and ∼30-day periods, we suspected a potential link to semi-lunar and lunar cycles.

The Moon follows three primary cycles that influence both moonlight intensity and gravitational forces on Earth (Fig. 2a). The synodic lunar cycle, lasting approximately 29.53 days, marks the transition between lunar phases – from new moon to full moon – as the Moon aligns with the Sun and Earth. This alignment also has gravitational effects, giving rise to spring tides around the new and full moons, and neap tides near the quarter moons, following a semi-lunar ∼14.8-day rhythm (Fig. 2b). The tropical (∼27.32 d) and draconic (∼27.21 d) months describe the Moon’s motion relative to the celestial equator and its orbital nodes, and together govern how high or low the Moon appears in the sky. The anomalistic cycle - averaging 27.55 days - traces the Moon’s path between its closest (perigee) and farthest (apogee) points in the elliptical orbit (Fig. 2c). This modulates the amplitude of spring tides. Since the synodic and anomalistic lunar cycles differ slightly in length, their alignment shifts over time. This periodic mismatch leads to occasional coincidences between full moons and new moons and perigee or apogee, termed as supermoons or micromoons.

**Fig. 2.**
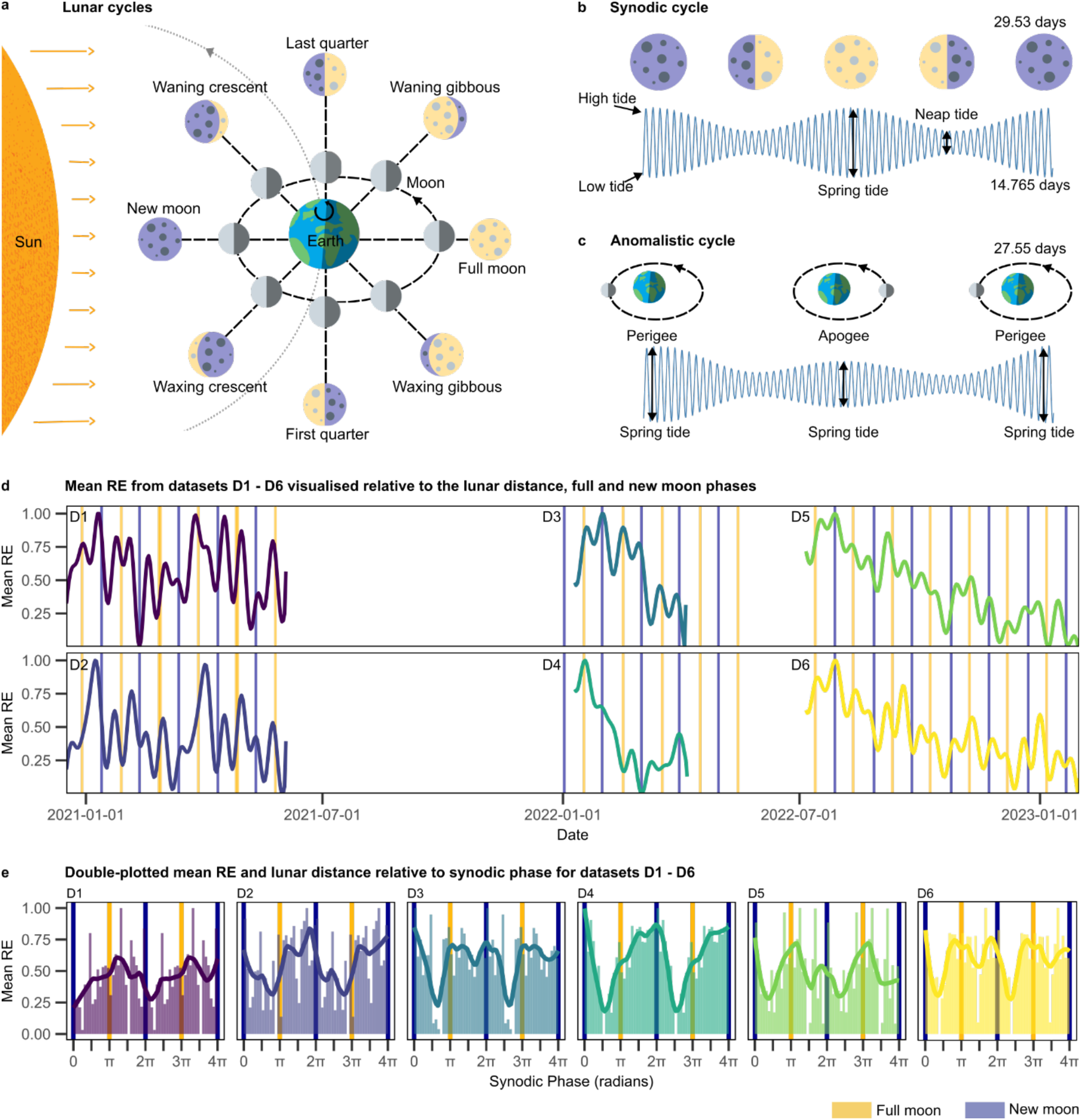
Evidence for semi-lunar periodicity in mean RE suggests that lunar cycles shape spontaneous behavioral fluctuations in mice under constant laboratory conditions. **(a)** Schematic overview of the Earth–Moon–Sun system, illustrating how the Moon’s orbit and rotation create distinct phases visible from Earth. While the Sun always illuminates half the Moon, the visible illuminated portion changes throughout the 29.53-day synodic month, resulting in full, new, and quarter moons. Sizes and distances not to scale. **(b)** The synodic lunar cycle generates periodic variation in lunar illumination and fraction visible from Earth. In parallel, the Moon’s gravitational influence produces tides with a 12.4-hour cycle. The alignment of Sun, Earth, and Moon during full and new moons leads to spring tides (every ∼14.77 days), while quarter moons produce weaker neap tides. **(c)** The Moon’s elliptical orbit creates the 27.55-day anomalistic cycle, during which its distance from Earth varies between perigee and apogee. Every 27.55 days, the Moon is close to Earth (in its perigee) and exerts maximal gravitational forces on Earth, resulting in high tidal amplitudes, whereas it exerts minimal gravitational forces when it is far from Earth (in its apogee). The synodic (∼29.53 days) and anomalistic (∼27.55 days) lunar cycles differ slightly in length. This mismatch causes the two cycles to drift in and out of phase over time. As a result, full and new moons sometimes coincide with perigee or apogee, producing supermoons and micromoons. **(d)** Mean RE (normalized from 0 to 1) from ColonyRack datasets D1–D6 plotted alongside full and new moon events, shows temporal alignment with full and new moon days suggestive of lunar modulation. **(e)** Double-plotted phase plots of mean RE relative to the synodic phase from D1–D6 show phase-locked peaks aligned with full and new moon phases in the lunar cycle. Vertical golden and dark blue bars mark full and new moon positions. Temporal and phase alignment metrics are reported in Table S4.

We examined RE time series in relation to two lunar cycles: the synodic cycle (∼29.5 days) and the anomalistic cycle (∼27.5 days). When RE values were plotted alongside normalized lunar illumination (scaled 0–1), we found a clear temporal alignment, with peaks in activity near full moon and new moon (Fig. 2d). Further analysis using double-plotted phase plots of RE relative to the synodic phase confirmed a much stronger and consistent alignment of behavioral peaks with full and new moon phases (Fig. 2e). Across all ColonyRack datasets, behavioral peaks showed a consistent temporal alignment with the lunar cycle and was quantified as the mean temporal offset between behavioral peaks and the nearest full or new moon (ΔDays), which was low (0.17 ± 0.94 days), indicating that activity peaks typically occurred within approximately one day of lunar events. The statistical significance of this alignment was assessed using a circular-shift test, in which the behavioral time series was systematically shifted with respect to the lunar cycle for 10,000 iterations while preserving the relative spacing between peaks. For each shift, temporal alignment with the lunar events was recomputed, generating a null distribution that retains intrinsic temporal structure but disrupts any fixed relationship to the lunar cycle. The observed temporal alignment was significantly stronger than expected under this null model (Fig. S3 a; Table S4). Phase relationships between behavior and lunar cycle were further characterized by converting temporal offsets into phase differences relative to the synodic cycle. Phase consistency across peaks was quantified using a circular measure of phase-locking value (PLV), defined as the mean resultant length of phase differences. PLV values were consistently high (mean PLV = 0.76 ± 0.10) with mean phase angles close to zero (0.05 ± 0.19 radians), indicating strong and consistent phase alignment. Statistical significance of phase locking was assessed using the same null model, with observed PLV values significantly greater than the null distribution. Together, these results demonstrate robust phase locking of behavioral rhythms to the synodic lunar cycle (Fig. S3 b; Table S4).

To further validate these findings, we conducted an independent experiment using running-wheel setups at two separate research facilities in Dresden, Germany. Over a period of 300 days, we recorded spontaneous locomotor activity from single- and pair-housed male and female C57BL/6 mice. The resulting four datasets – R1 (Facility 1) and R2 (Facility 2), each comprising male and female groups – revealed consistent semi-lunar rhythmicity and additional infradian harmonics (Fig. 3a, b). Power spectral density analysis (Fig 3c) and spectrograms (Fig. S4) confirmed stable power clustering around these infradian periods. When activity was plotted against the synodic phase, double-plotted phase diagrams showed clear and consistent alignment of behavioral peaks with both full and new moon (Fig. 3d). The mean temporal offset from the nearest full or new moon was negative but within a day (ΔDays = −0.21 ± 0.34 days) and statistically significant when compared against a null distribution generated by circularly shifting the behavioral time series relative to lunar cycle (Fig. S5 a; Table S4). Phase-locking values were consistently high (mean PLV = 0.66 ± 0.08) with mean phase angles near zero (−0.08 ± 0.08 radians) and statistically significant against a null distribution (Fig. S5 b; Table S4). Together these results confirm that mouse behavior, even under stable and controlled environmental conditions, contains rhythmic components that align with full and new moon phases of the lunar cycle.

**Fig. 3.**
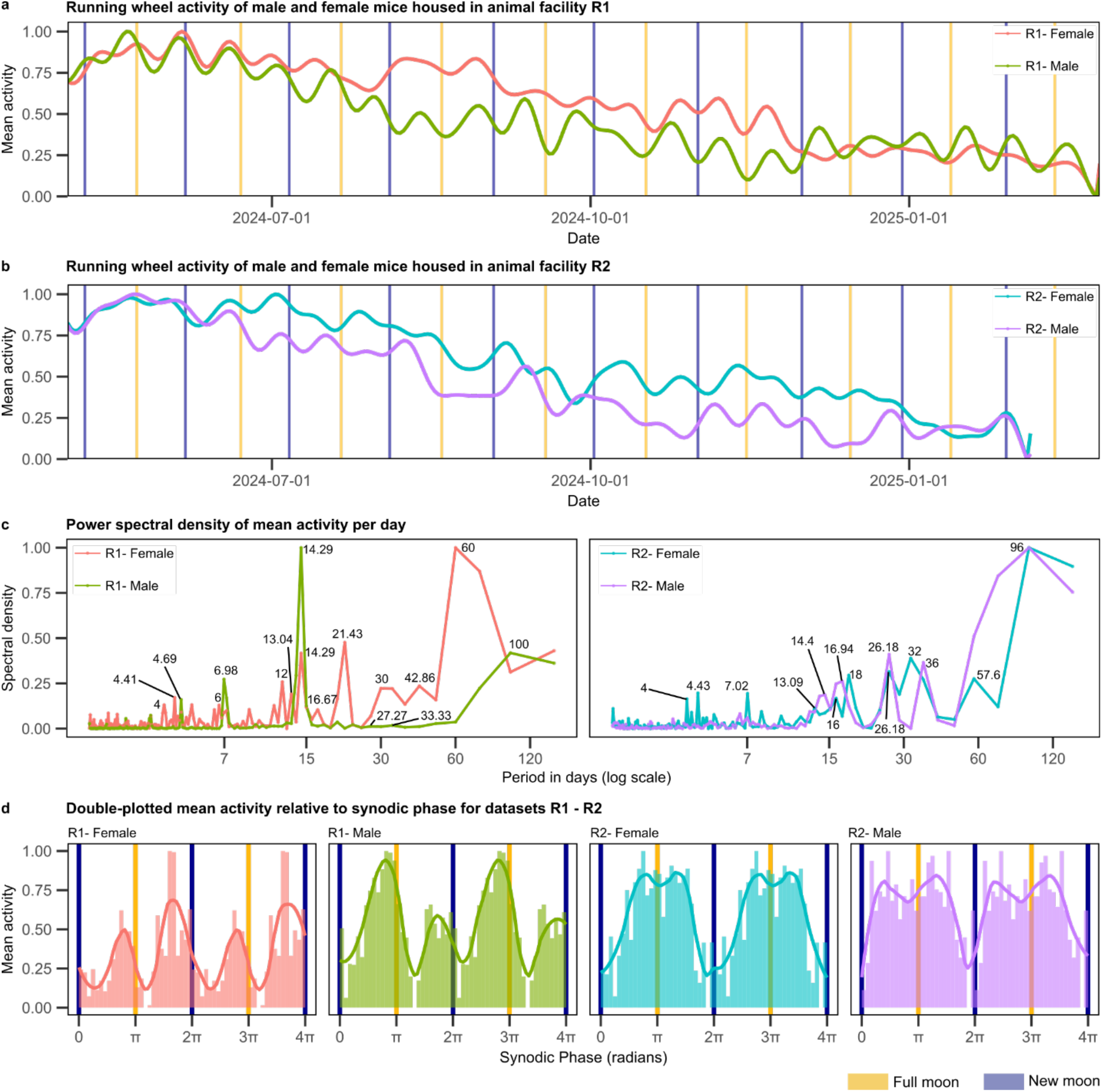
Semi-lunar and lunar periodicity in running wheel activity recorded in different animal facilities in Dresden. **(a, b)** Running wheel activity (normalized from 0 to 1) from male and female mice in two separate animal facilities (R1 and R2) also shows infradian rhythmicity. **(c)** Power spectral analysis (normalized from 0 to 1) shows clustering of power at ∼15-day intervals, consistent with semi-lunar periodicity. Numbers in the plot mark the peaks in days. **(d)** Double-plotted phase plots of mean activity relative to the synodic phase from R1–R2 show phase-locked peaks aligned with full and new moon phases in the lunar cycle. Vertical golden and dark blue bars mark full and new moon. Temporal and phase alignment metrics are in Table S4.

Similar to how the anomalistic cycle influences the amplitude of semi-lunar tides, it appears to modulate the amplitude of behavioral fluctuations. Specifically, when the moon is at apogee, its farthest point from Earth, we observed dips in activity across datasets, unless this coincided with the full or new moon (Fig. S6).

### Semi-lunar behavioral rhythms are conserved across independent long-term mouse datasets

To confirm that these semi-lunar rhythms are not specific to or even artifacts of our laboratory conditions, we reanalyzed data from a study by Acosta-Rodriguez et al., 2022 ^27^. The dataset recorded in Dallas (USA) included running wheel activity of single-housed male C57BL/6J mice over their entire lifespan (max 1308 days), starting at 8 weeks of age. All mice had unrestricted access to food for six weeks and then were randomly divided into six different groups with food access: *ad libitum* (AL) or under caloric restriction (CR, 70% of AL intake) and varying food availability. For visualization and analysis, a subset of the data was selected covering the period from 12 Sep 2017 to 8 Feb 2021 (1246 days) which included 63 mice (AL: n=5, CR-night-12h: n=12, CR-night-2h: n=11, Cr-day-12h: n=11, CR-day-2h: n=12 and CR-spread: n=12). Importantly, all animals were maintained on a purified diet with a consistent nutritional composition throughout the entire experiment. This is critical because standard grain-based vivarium diets can vary across batches due to seasonal changes in grain sources, potentially introducing food-driven seasonal rhythms. Using a purified diet eliminates this source of variability.

All groups maintained a normal dark-time locomotor activity pattern throughout life, except for the day-fed (food at day-time) and CR-spread mice (continuous food availability), which showed increased daytime activity. Longitudinal plots revealed slow-oscillatory patterns in daily running activity (Fig. 4a). The spectral analysis confirmed the presence of semi-lunar rhythmicity and additional infradian harmonics (Fig. 4b, Fig. S8). Double-plotted phase plots of mean activity relative to the synodic phase showed consistent alignment of behavioral peaks with full and new moon phases (Fig. 4c). The average temporal offset from the nearest lunar event was again negative but small (ΔDays = −0.66 ± 0.78) and statistically significant relative to a null distribution generated by circularly shifting the behavioral time series relative to lunar cycle (Fig. S9 a; Table S4). Phase consistency was likewise high (mean PLV = 0.74 ± 0.04), with mean phase angles close to zero (−0.16 ± 0.17 radians), and significantly exceeded the null expectation (Fig. S9 b; Table S4). This confirmed the reproducibility of our observation.

**Fig. 4.**
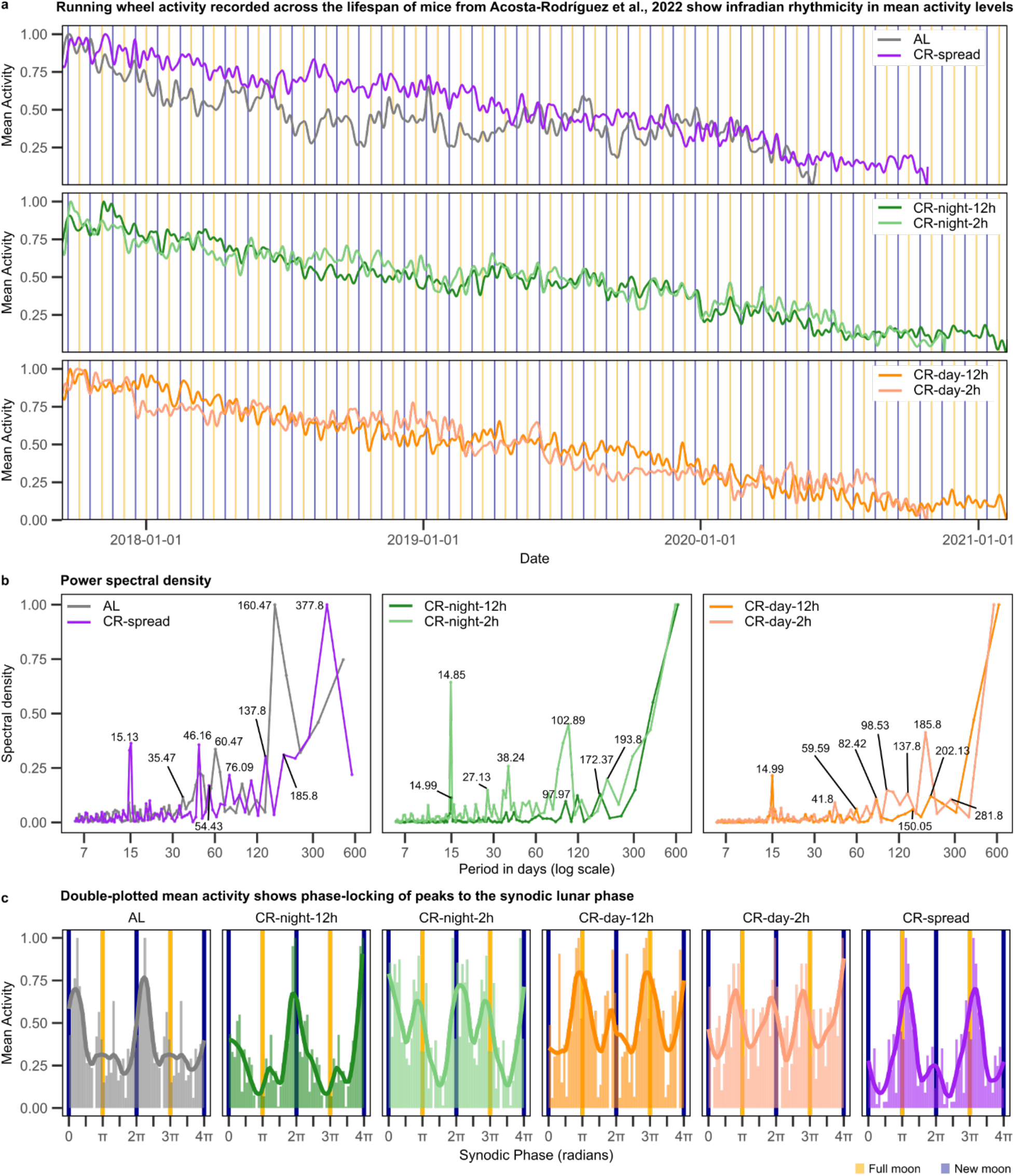
Semi-lunar and lunar periodicity in running wheel activity of mice across their lifespan from an independent study - Acosta-Rodriguez et al., 2022 reveals conserved infradian patterns in mouse behavior. **(a)** Group-averaged time series of daily running-wheel activity (normalized from 0 to 1) for AL (n=5), CR-night-12h (n=12), CR-night-2h (n=11), CR-day-12h, (n=11), CR-day-2h (n=12) and CR-spread mice (n=12) mice from 12 Sep 2017 to 8 Feb 2021 (1246 days) show patterns of slow, rhythmic fluctuations under constant laboratory conditions. **(b)** Spectral analysis (normalized from 0 to 1) of daily activity time series reveals consistent clustering of power at ∼15- and ∼30-day periods, corresponding to semi-lunar and lunar periodicities. Numbers in the plot mark the peaks in days. **(c)** Double-plotted phase plots of mean activity relative to the synodic phase show phase-locked temporal alignment of behavioral peaks with full and new moon phases in the lunar cycle. Vertical golden and dark blue bars mark full and new moon. Temporal and phase alignment metrics are reported in Table S4.

### Persistence in the absence of a circadian clock

To investigate whether lunar-coupled rhythms depend on circadian mechanisms, we reanalyzed data from a study by Izumo et al. (2014) ^28^. The dataset included *Bmal1* conditional knockout (cKO) mice, which lack a functional superchiasmatic nucleus (SCN) due to forebrain-specific deletion of the *Bmal1* gene, and appropriate controls. The mice were sequentially exposed to a series of lighting conditions: a standard light–dark (LD) cycle, constant darkness (DD), constant light (LL), and finally a return to LD. The study showed that these mice are behaviorally arrhythmic under constant dark or light conditions (DD and LL), providing a powerful model to assess infradian rhythmicity in the absence of a central circadian oscillator.

Despite lacking a functional SCN, *Bmal1* cKO mice exhibited clear infradian oscillations in mean activity levels (Fig. 5a). Spectral analysis of daily activity time series showed strong power at semi-lunar periods and lunar periods (Fig. 5b, Fig. S10). Phase alignment to the synodic lunar cycle remained intact even in the cKO group (Fig. 5c). The average offset from the nearest lunar event was again negative but small (ΔDays = −0.15 ± 0.39) and significant relative to a null distribution (Fig. S11 a; Table S4). Phase-locking values were consistently high (mean PLV = 0.77 ± 0.03) with mean phase angle close to zero (−0.04 ± 0.09 radians) and significantly exceeded the null expectation (Fig. S11 b; Table S4).

**Fig. 5.**
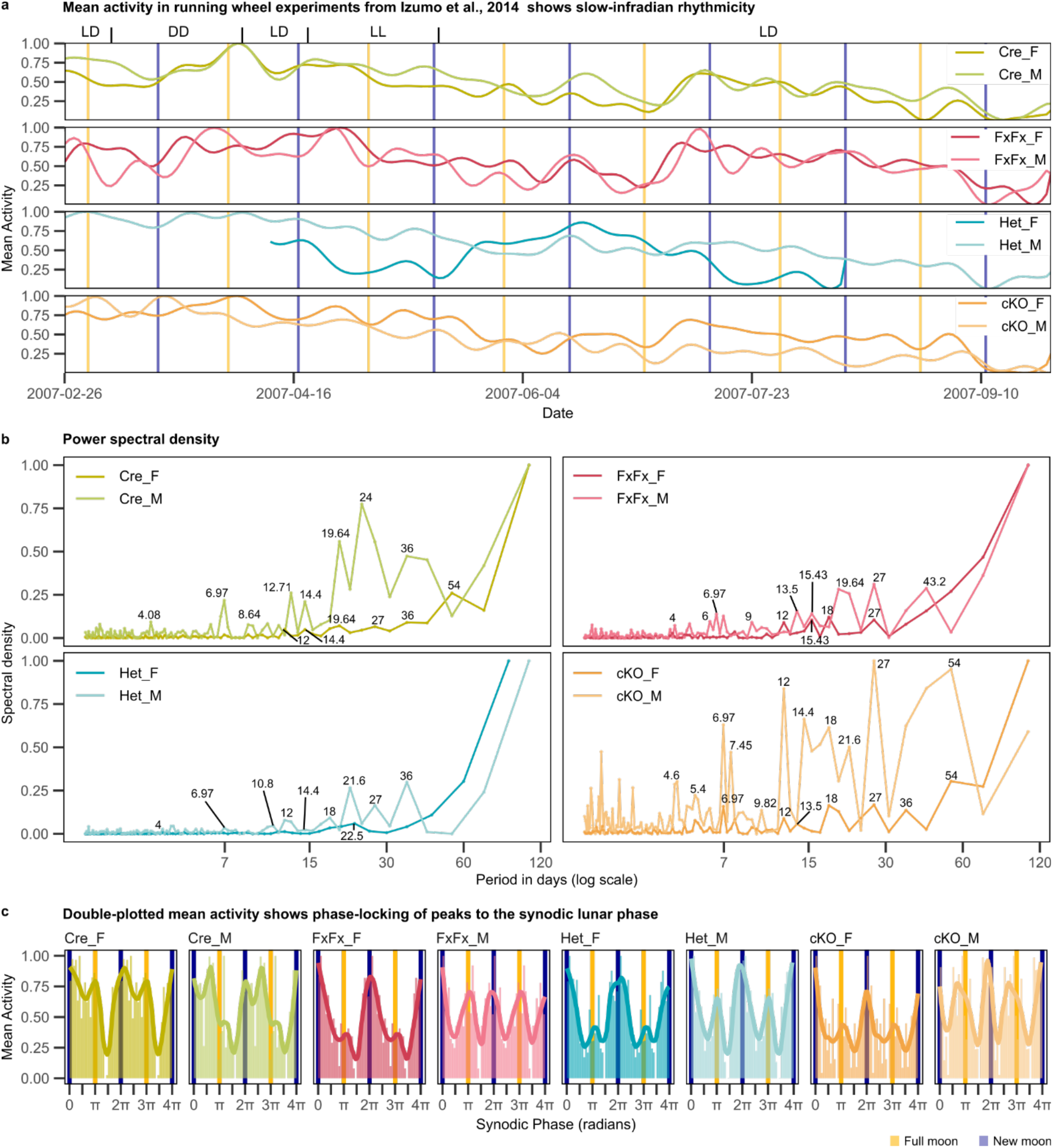
Running wheel data from an independent study - Izumo et al., 2014 confirms the presence of semi-lunar and lunar periodicity despite the brain knockout of *Bmal1*. **(a)** Time series of daily running wheel activity (normalized from 0 to 1) for Cre (*CamiCre*+, female, n=8; male, n=5), FxFx (*Bmal1*^fx/fx^, female, n=9; male, n=9), Het (*CamiCre+; Bmal1*^fx/+^, female, n=8; male, n=8), and cKO (*CamiCre+; Bmal1*^fx/fx^, female, n=8; male, n=10) mice show infradian rhythmic fluctuations despite larger fluctuations in response to changes in light-dark condition (marked on top) and genotypes. Mice were initially housed under a 12:12 hour light-dark cycle (LD), followed by 4 weeks in constant darkness (DD), 2 weeks back in LD, 4 weeks in constant light (LL), and finally a return to LD. **(b)** Power spectral density plots (normalized from 0 to 1) show clustering of power at ∼15 to ∼30-day periods. Numbers in the plot mark the peaks in days. **(c)** Mean activity values plotted against the synodic lunar phase demonstrate phase-locked alignment of behavioral peaks with specific positions in the lunar cycle. Vertical golden and dark blue bars mark full and new moon. Temporal and phase alignment metrics are reported in Table S4.

These results show that lunar-coupled behavioral rhythms are not dependent on the central circadian clock. Instead, they may emerge from alternative mechanisms that can operate independently of the SCN, potentially through sensitivity to environmental cues beyond light.

### Behavior and geomagnetic fluctuations show temporal coupling at shared frequencies

The observation of behavior rhythms at periods close to principal lunar cycles and their harmonics hinted at their lunar origin. Lunar gravitation drives tides in large water bodies but not in small fluid systems such as lakes, coffee cups, or cells, raising the question of how lunar cycles could robustly influence murine physiology and behavior. We hypothesize that an intermediate geophysical factor, modulated by the Moon’s orbital motion and Sun’s rotation, mediates this effect.

Geomagnetic activity contains both lunar tidal and solar-rotation related periodic components. Amplitude spectra of observed geomagnetic field time series show clear quasi-monthly oscillations centered around ∼27-28 days period, originating in the magnetosphere and ionosphere ^29^, reflecting the effect of the solar synodic rotation period (Carrington rotation) of ∼27 days ^30,31^ and lunar cycles ^32,33^. In addition, quasi-fortnightly oscillations near 14 days are consistent with harmonics and sidebands of the solar synodic (∼27 d) and lunar synodic (∼29.5 d) cycles^29^. Despite their very close periods, lunar and solar oscillations arise from distinct physical mechanisms and are generally not phase-locked. The superposition of these harmonic oscillations produces a time-varying interference pattern that can modulate the net geomagnetic signal experienced by organisms.

To test whether geomagnetic fluctuations covary with behavior, we analyzed geomagnetic field data from the closest observatories alongside mouse activity. Across all datasets, behavioral peaks closely followed geomagnetic peaks (Fig. 6a, b; Fig. S12, S14, S16, S18). The mean temporal offset was small (ΔDays = 0.54 ± 1.04), with peak activity typically occurring within a day of a geomagnetic peak. The statistical significance of this alignment was assessed against a null distribution, in which the behavioral time series was circularly shifted with respect to the geomagnetic time series for 10,000 iterations. The observed temporal alignment was significantly stronger than expected under this null model (Fig. S13 a, S15 a, S17 a, S19 a; Table S5). Phase-locking values were high (mean PLV = 0.75 ± 0.08), with mean phase angles close to zero (0.12 ± 0.25 radians; Table S5). Statistical significance of phase locking was assessed using the same null model, with observed PLV values significantly greater than the null distribution (Fig. S13 b, S15 b, S17 b, S19 b; Table S5). indicating robust synchronization unlikely to occur by chance. This suggests a phase-lagged relationship, in which changes in environmental electromagnetic field precede, and potentially influence, behavioral rhythms.

**Fig. 6.**
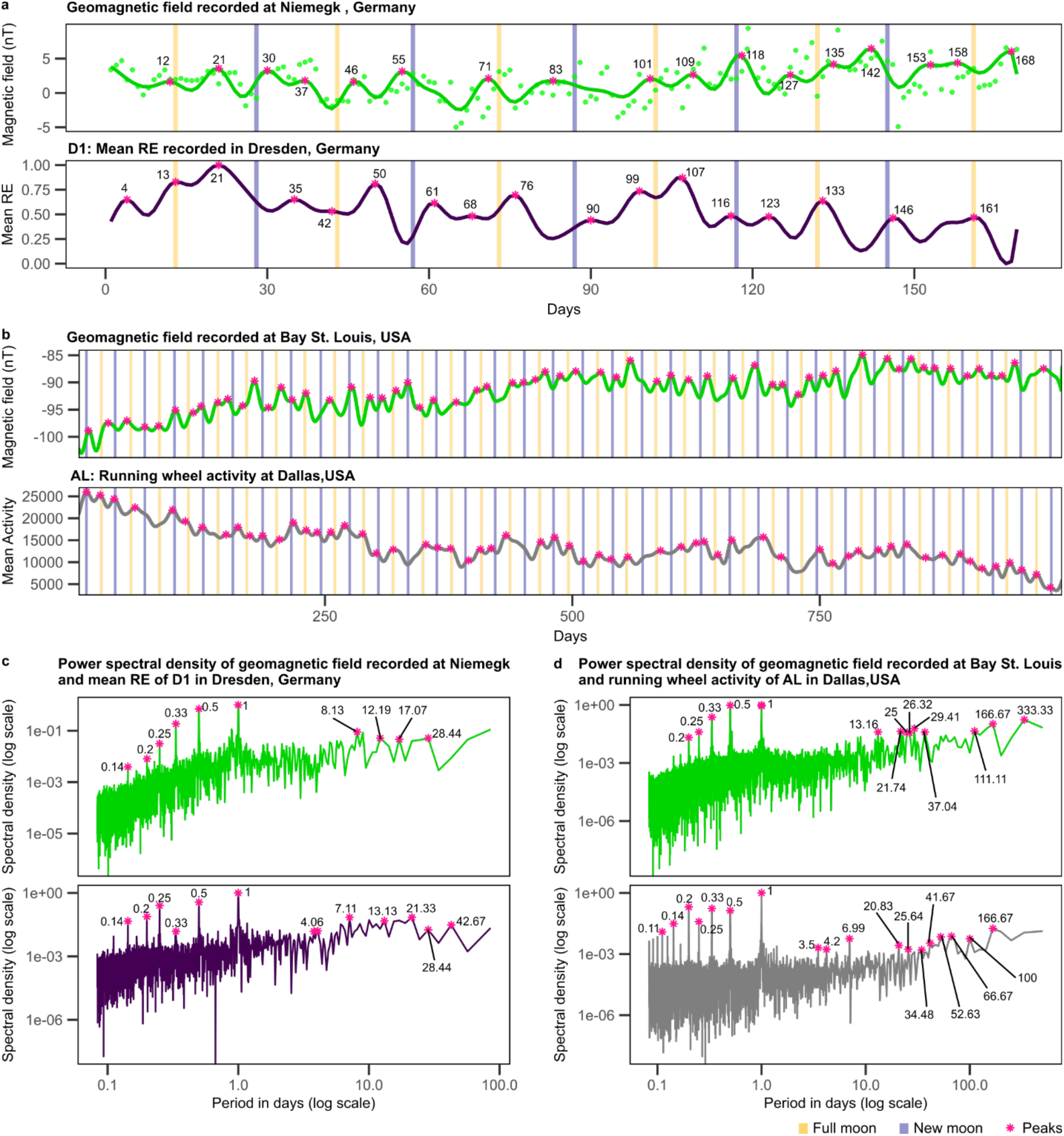
Local geomagnetic field variations and mouse behavioral activity co-vary with overlapping dominant frequencies. **(a)** Geomagnetic field variations recorded at Niemegk observatory, Germany, and mean RE from dataset D1 (App^NL-G-F/NL-G-F^, n = 33) recorded at DZNE, Dresden, from 17 Dec 2020 to 31 May 2021 (166 days). **(b)** Geomagnetic field variations recorded at Bay St. Louis observatory (MS), USA, and running wheel activity from dataset AL (n = 5; Acosta-Rodriguez et al., 2022) recorded at Dallas (TX), USA, from 12 Sep 2017 to 8 Feb 2021 (1246 days). **(c)** Power spectral density for geomagnetic field recorded at Niemegk observatory (top) and mean RE from ColonyRack dataset D1 (bottom). **(d)** Power spectral density for geomagnetic field recorded Bay St. Louis (top) and running wheel activity of dataset AL (bottom). Numbers on the time series and power spectra mark the peaks in days. Temporal alignment, phase locking values, and corresponding statistics are reported in Table S5. For all data sets, the North-directed component of the geomagnetic field is shown.

Spectral analysis revealed overlapping dominant frequencies in both signals. In addition to synodic and semi-synodic rhythms, shared oscillations occurred at diurnal periods (Fig. 6c, d). In geomagnetic spectra, strong peaks at 1–4 cycles per day (harmonics at 24, 12, 8, and 6 hour) were observed in both magnetospheric and ionospheric components, with the ionospheric signal showing higher signal power ^29^. These correspond to the solar daily variation and its sub-harmonics, reflecting activity in day-side ionospheric current systems ^34^. Superimposed on these were weaker peaks at nearby periods consistent with lunar daily variations (harmonics at 25.7, 12.4, 8.2, and 6.1 hours), which follow Chapman’s phase law and can be described as harmonic oscillations governed jointly by solar time and the Moon’s position relative to the Sun ^32^.

To examine the association between geomagnetic fluctuations and behavioral activity in the ultradian range, we plotted the local geomagnetic field and behavioral activity across a 48-h double-plotted window. Consistent with the spectral structure of the geomagnetic field, local geomagnetic activity had two recurring peaks within each 24-h cycle: a smaller peak occurring in the early morning (3:00 to 6:00 hours) and a larger fluctuation around midday to early afternoon (11:00 to 13:00 hours) (Fig. 7 a, b, c, top panel). These fluctuations correspond to the solar daily variation and its harmonics generated by day-side ionospheric current systems, with weaker contributions from nearby lunar daily harmonics.

**Fig. 7.**
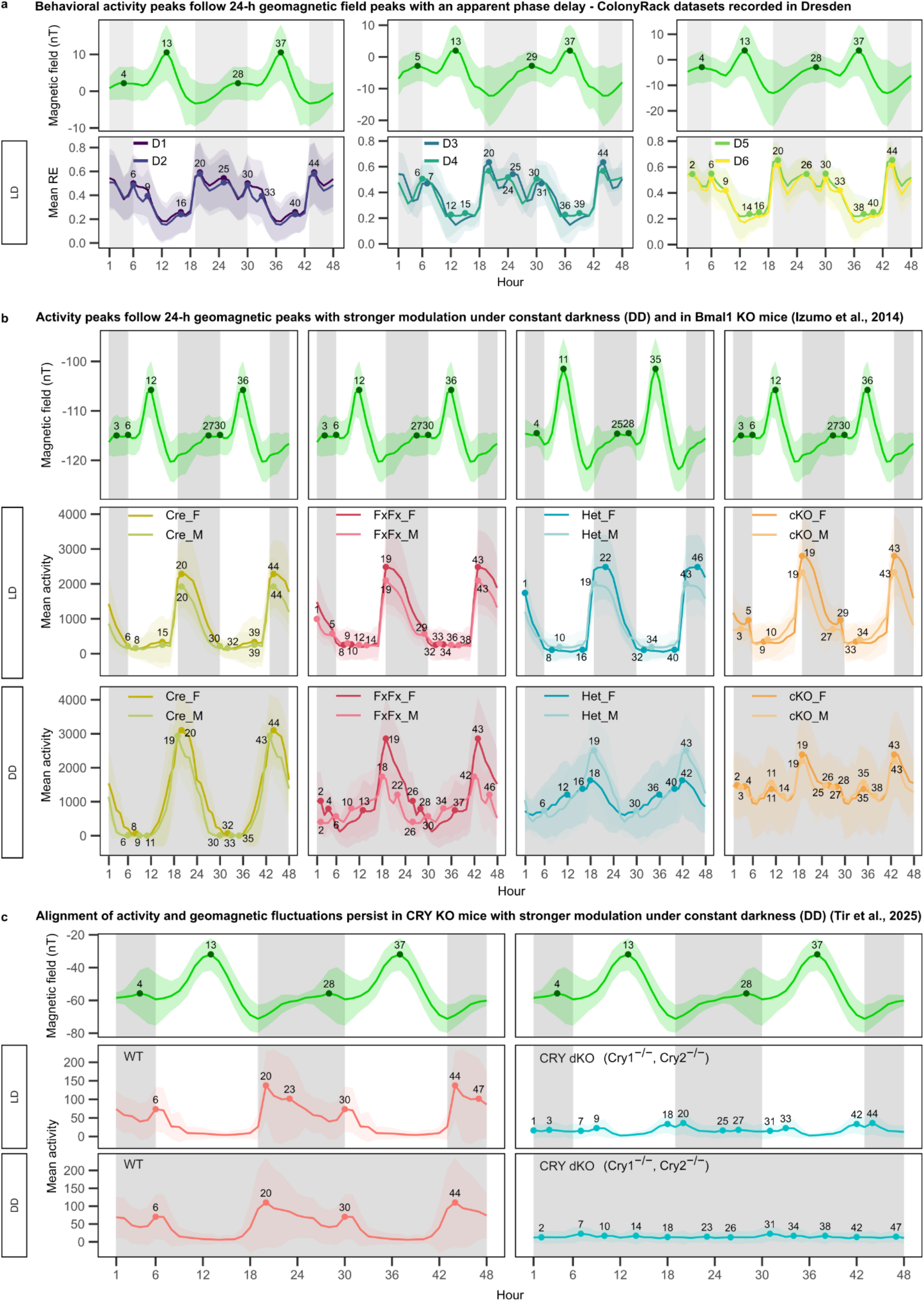
Geomagnetic–behavioral alignment in the 24-h range. **(a)** Local geomagnetic field variation recorded at the Niemegk observatory and mouse exploratory activity (mean RE) recorded in the ColonyRack system in Dresden, plotted across a 48-h double-plotted window to visualize alignment in the daily range. **(b)** Local geomagnetic field variation recorded at Fredericksburg, USA, and mouse running-wheel activity recorded in Evanston, USA, plotted across a 48-h double-plotted window under light–dark (LD) and constant darkness (DD) conditions. Data are shown for Cre controls (*CamiCre*+, female, n=8; male, n=5), FxFx controls (*Bmal1*^fx/fx^, female, n=9; male, n=9), Het controls (*CamiCre+; Bmal1*^fx/+^, female, n=8; male, n=8), and *Bmal1* conditional knockout mice (cKO; *CamiCre+; Bmal1*^fx/fx^, female, n=8; male, n=10). **(c)** Local geomagnetic field variation recorded at Hartland, UK, and mouse home-cage activity recorded using Digital Ventilated Cage (DVC) monitoring in Oxford, UK, plotted across a 48-h double-plotted window under LD and DD conditions. DVC activity was recorded from C57BL/6J mice (WT; n = 6) and cryptochrome double knockout mice (*Cry1*^−/−^, *Cry2*^−/−^; CRY dKO; n = 6). Grey shading indicates the dark phase. Lines show mean activity and ribbons indicate standard deviation. Detected maxima are marked by points and labelled according to their timing within the 48-h window.

Under light–dark conditions, behavioral activity was strongly shaped by light transitions, with prominent peaks occurring around lights-on and lights-off. In the double-plotted representation, these light-driven activity maxima were evident at approximately 06:00, 20:00, 30:00, and 44:00 (Fig. 7 a, b, c, second panel). However, beyond these expected responses to the light schedule, additional activity peaks can be observed near the geomagnetic maxima, typically following them with a short lag of approximately 0–2 h.

The relationship is more apparent under constant darkness (DD), where behavior is no longer directly driven by light transitions although some residual temporal structure likely reflects prior entrainment to the light cycle. This effect was particularly evident in *Bmal1* conditional knockout mice, which lack circadian rhythmicity (Fig. 7 b, third panel). This suggests that, when the circadian clock is disrupted and light cues are absent, behavioral activity may become more sensitive to geomagnetic fluctuations.

In mammals, cryptochromes, particularly CRY2, have been proposed as a candidate magnetically sensitive protein. However, much of the supporting evidence comes from heterologous expression studies in insects rather than from direct behavioral evidence in mammals^35–37^. Given this proposed link between CRY2 and magnetic sensitivity, we examined whether geomagnetic–behavioral alignment persists in mice lacking both major mammalian cryptochromes.

For this analysis, we used data made available by Tir et al.^38^. Using Digital Ventilated Cage (DVC) system in combination with capacitive sensors mounted under individual cages locomotor activity was recorded from C57BL/6J mice (WT; n = 6) and cryptochrome double knockout mice (*Cry1*^−/−^, *Cry2*^−/−^; CRY dKO; n = 6) for 19 days with LD conditions for first 8 days and DD for the rest. Under LD conditions, CRY dKO mice showed light-dependent activity patterns, as expected, but also displayed activity maxima near geomagnetic field fluctuations (Fig. 7 c). Under DD conditions, when activity was no longer directly shaped by light cues, activity maxima continued to occur close to geomagnetic peaks. This suggests that the observed diurnal geomagnetic modulation is not abolished by loss of CRY1/2 function. Interpretation of these data must remain cautious, however, because CRY1/2 knockout mice exhibited very low overall activity levels, resulting in comparatively flat activity peaks.

The temporal alignment of behavioral rhythms with geomagnetic fluctuations both in the ultradian and infradian range provides a plausible explanation for the origin of these periodic structures in the spontaneous activity of mice, suggesting that geomagnetic field may act as an environmental stimulus mediating the combined influence of lunar cycles and solar rotation on physiology and behavior.

## Discussion

Our findings reveal a previously unrecognized dimension of behavioral organization in mammals: a rhythmic structure driven not only by the circadian day-night cycle but also by longer infradian oscillations possibly tied to the solar-lunar interaction with the Earth’s geomagnetic field. Across multiple spatially and temporally independent cohorts, and experimental conditions, we consistently observed rhythms in both exploratory and spontaneous locomotor activity around the solar and lunar synodic (∼27 ± 3 days) and semi-synodic (∼ 14 ± 1 days) periods. Across all datasets, behavioral peaks showed a consistent temporal alignment with the lunar cycle. The average temporal offset from the nearest full or new Moon was small (ΔDays ≈ –0.25 ± 0.75), indicating that peak activity tended to occur within about a day of a lunar phase transition. Phase-locking values were generally high (mean PLV = 0.74 ± 0.07), with mean phase angle near zero (−0.07 ± 0.17). Temporal offset and PLV were significant relative to a null distribution generated by circularly shifting the behavioral time series, indicating that the observed relationship was unlikely to arise by chance. Rayleigh tests further confirmed that behavioral peaks were not uniformly distributed across the synodic lunar cycle. We tested both unimodal and bimodal alternatives to distinguish clustering around a single preferred lunar phase from axial clustering around two opposing phases. The significant bimodal Rayleigh test supports a semi-lunar phase structure, indicating that activity peaks aligned with both full and new moon phases rather than occurring randomly across the lunar month (Table S3).

The consistency of this pattern across different continents and years shows that the observed phase alignment is a robust and reproducible feature of mouse behavior, not a coincidental or batch-specific effect. Our reanalysis of *Bmal1* conditional knockout cKO) mouse data from Izumo et al. ^28^ provides critical support for this interpretation. Despite the loss of a functional SCN and the absence of circadian behavior under constant conditions, *Bmal1* cKO mice still exhibited robust infradian oscillations, including semi-synodic rhythmicity, in overall activity levels. Spectral analyses revealed power at semi-synodic periods, and phase-locking to lunar cycles was preserved. These observations suggest that the well-established molecular clock in the SCN underlying mammalian circadian oscillations is not necessary for the generation or maintenance of these infradian rhythms.

Fluctuations in the geomagnetic field on the ground contain multiple time harmonics with periods around ∼27 ± 3 and ∼ 14 ± 1 days. These magnetic fluctuations are global-scale and originate mainly in the magnetosphere and ionosphere, which are modulated by solar rotation and lunar orbital period ^29^. In the diurnal or circadian range, geomagnetic spectra contain peaks at 1–4 cycles per day (∼ 24, 12, 8, and 6 hour) attributed both to solar and lunar daily variations ^39,33^. At periods longer than the synodic period, spectral peaks are present at ∼54, 81, 135, 162, and 243 days. These spectral peaks are multiples of the solar rotation period and associated with solar activity variations ^40^.

The evidence of a temporal alignment of behavioral rhythms with geomagnetic fluctuations is multifold. First, the Fourier spectra of both the rhythms measured across multiple location and time periods contain overlapping spectral components. This is evident not only in the synodic and semi-synodic periodbands, but also in the diurnal range at ∼ 24, 12, 8, and 6 hours. Regarding longer periods, we consistently observed periods that are multiples of the solar rotation, e.g. close to ∼ 54, 81, and 135 days. Second, in the infradian range, peaks in the geomagnetic fluctuations seem to be aligned with the behavioral peaks with an offset (ΔDays = 0.84 ± 1.32) and the behavioral peaks following the geomagnetic peaks. Across all datasets collected from different locations worldwide and spanning nearly two decades (2007–2025), we observed a significant phase-locking value (mean PLV = 0.75 ± 0.08) with mean phase angles close to zero (0.14 ± 0.25 radians). Both the temporal offset and PLV differed significantly from a null distribution generated by circularly shifting the time series, indicating that the observed alignment was unlikely to result from chance. Third, in the diurnal and ultradian range, behavioral activity showed fluctuations that corresponded to geomagnetic maxima with a small lag of approximately 0–2 hours. This alignment was more pronounced under complete darkness and in mice with impaired circadian clock function, suggesting that geomagnetic modulation of behavior may become more detectable when dominant light-driven or endogenous circadian signals are reduced. The recurring lag between geomagnetic and behavioral peaks points to a plausible delayed sensory responsiveness to geophysical cues, suggesting a phase-lagged relationship, in which changes in environmental geomagnetic conditions may precede and potentially modulate behavioral timing. This is consistent with previous reports that responses to magnetic-field changes can be delayed, possibly reflecting the time needed to integrate a weak geophysical signal and translate it into behavioral output^41,42^. Geomagnetic field may act as an environmental stimulus mediating the combined influence of lunar and solar periods on physiology and behavior on Earth.

All life on Earth evolved under the constant presence of the geomagnetic field. Its omnipresence makes it unsurprising that many animal species have evolved ways to sense it and use it for navigating their world. The Earth’s magnetic field offers rich, map-like information: polarity distinguishes normal from reversed field directions, inclination gives the tilt of the field lines relative to the horizontal, declination is the angle between magnetic and geographic north, and intensity is the strength (magnitude) of the field. For example, the European robin, a well-studied long-distance migrant, relies on a magnetic inclination compass during its seasonal journeys ^43^. Magnetoreception, including compass and in some cases map-like use of the geomagnetic field, has also been demonstrated in sea turtles, fishes like eel, salmon and bonnethead sharks, and amphibians like newts ^41^. Even some mammals exhibit this ability: subterranean mole-rats use the magnetic field to orient themselves through complex underground burrow systems ^44^.

On top of this steady presence, the Sun and Moon create regular, periodic fluctuations in the Earth’s electromagnetic environment. Every organism on Earth has also been exposed to these invisible shifting in environmental cues, so that animals might also have evolved mechanisms to sense them and adjust the timing of their behaviors. Any behavioral rhythms influenced by geomagnetic cues would therefore experience a multi-periodic, sometimes reinforcing, sometimes opposing combination of lunar and solar-driven variations, with solar-driven geomagnetic variations typically having a larger amplitude ^29^.

The Moon is a strong, yet enigmatic zeitgeber, whose influence spans illumination (synodic cycle), gravitational variation (synodic and anomalistic cycle), and tide-generation (diurnal and fortnightly harmonics). In marine and amphibious species, moonlight and gravitational tides are known to entrain rhythms in reproduction, locomotion, and feeding ^17,18^. While such lunar synchronization mechanisms are increasingly understood in marine species, their existence and functional relevance in terrestrial mammals, including humans, has always remained very controversial ^45^. Decades of research have searched for lunar correlations with menstrual cycles ^46,47^, sleep patterns^48,49^, and mood disorders, including suicides ^50,51^, yielding inconsistent and often inconclusive results. However, recent longitudinal studies suggest that lunar associations may become detectable when analyzed at the appropriate temporal and individual scale. Menstrual cycles have been reported to transiently synchronize with the synodic lunar cycle, and in some records also with tropical and anomalistic lunar cycles^19^. Similarly, mood rhythms in patients with rapid cycling bipolar disorder have been shown to align with the 14.8-day spring–neap cycle, the 13.7-day declination cycle and the perigee-syzygies (‘supermoons’)^52^ indicating phase-locking not only to the synodic cycle but also to other lunar orbital components. Sleep timing has also been reported to vary across the lunar month, with a recent human wrist-actimetry study showing that sleep begins later and is shorter on nights preceding the full moon across Indigenous Toba/Qom communities in rural Argentina with and without access to electricity, as well as in a highly urbanized population in the United States, representing populations exposed to or relying on different levels of lunar illumination information^53^. Our findings thus place mouse exploratory activity within a growing literature reporting lunar or semi-lunar organization of mammalian physiology and behavior. They further suggest that these earlier observations may need to be reevaluated in light of a possible sensitivity of terrestrial mammals ^54^, possibly including humans ^55,56^, to solar–lunar–driven geomagnetic fluctuations, which could provide a unifying mechanism that underlies some of the reported associations between behavior and physiology and lunar phase.

We found that both female and male mice displayed synodic and semi-synodic rhythmicity, indicating that these cycles are unlikely to arise from the female estrous rhythm (4-5 days)^57^. Interestingly, males showed higher oscillatory amplitude, hinting at possible sex-specific sensitivity to infradian modulation, an observation that warrants further exploration, particularly in relation to hormonal cycles and their potential interaction with solar-lunar geomagnetic variations.

Might our findings be more simply explained by direct gravitational effects? We consider this unlikely on biophysical grounds. The gravitational acceleration of the Moon at the surface of the Earth is about 1 to 3 millionth of 1 g ^58^, with the tidal acceleration being an order of magnitude smaller. The force exerted onto a mammalian cell by this gravitational pull is approximately 0.03 fN. For Piezo1, the best studied mechanosensitive channel, the force to activate gating is estimated to be 4.7 +/- 0.3 nN/m ^59^. For a channel of 5nm radius and a membrane thickness of 5nm this converts to approximately 10 to 100 pN, which is 10^6^ to 10^9^ times greater than the lunar gravitational effect. The solar tidal acceleration, for comparison, is about half the Moon’s (the Sun is much larger, but also further away). A recent study analyzing longitudinal actigraphy recordings in humans and captive titi monkeys, in relation to lunar cycles and estimated lunisolar gravimetric variation reported that sleep timing is synchronized not primarily with moonlight, but with semi-lunar and lunar gravimetric force maxima that occur around full and new moon^60^. This is consistent with our observation that high behavioral activity tends to align with full and new moon phases. The relative positions of Sun-Moon-Earth at full and new moon phases is not only associated with maxima in lunisolar gravimetric forces, but also with changes in the geomagnetic field. Consequently, we believe that a sense of the impact of Sun and Moon on mouse behavior is more likely to be mediated through geomagnetic fluctuations as an environmental intermediate rather than through a direct mechanical sense for gravitational forces. But future research should remain open to this theoretical possibility as well.

How mice might detect such weak geomagnetic variation (in the nT range) remains unresolved, but two broad mechanisms are most relevant. The first is a light-dependent radical-pair mechanism, most often discussed in relation to retinal cryptochromes^54,61,62^. In birds, CRY4 is considered a strong candidate magnetoreceptor, but mammals do not have CRY4 and instead express CRY1 and CRY2, which are primarily known for their roles in the circadian clock^63,64^. Nevertheless, mammalian cryptochromes remain of interest: human CRY2 can support light-dependent magnetic responses in Drosophila, and weak electromagnetic fields have been reported to alter intracellular ROS in a CRY1/CRY2-dependent manner in human cells^36,65^. Thus, CRY-dependent magnetosensitivity in mammals is possible, but remains mechanistically unresolved^64^. In our analysis of CRY1/2 double-knockout mice, activity peaks still showed some correspondence with diurnal geomagnetic maxima, particularly under complete darkness. However, overall activity in these mice was very low and the resulting activity peaks were relatively flat, limiting any interpretation. Thus, while these data suggest that diurnal behavior–geomagnetic alignment is not completely abolished in the absence of CRY1/2, future experiments will be required to determine how and if CRY2 contributes to mammalian magnetoreception.

A second possibility is a magnetite- or biogenic magnetic-particle-based mechanism, in which magnetic particles could transduce geomagnetic variation into mechanical or cellular signals. This mechanism has been considered particularly relevant in mammals with robust magnetic orientation, such as bats^66^ and mole-rats^67,68^. Evidence for magnetic sensitivity also exists in surface-dwelling rodents: wood mice orient nests relative to the magnetic field, and this orientation is disrupted by weak radiofrequency fields, consistent with a radical-pair-like contribution^69,62^. However, direct anatomical evidence for a magnetite-based receptor in mice is still limited. Our data therefore do not distinguish between cryptochrome-dependent, magnetic-particle-based or hybrid mechanisms, but they suggest that mouse behavior is sensitive to natural geomagnetic variation and motivate future experiments targeting the underlying sensory pathway.

Taken together, our results support the existence of a long-period oscillation in mammals - one that encodes celestial periodicities independently of the traditional molecular feedback loops of the circadian clock. Whether this is an endogenous rhythm or physiological sensitivity to geomagnetic fluctuations remains an open and exciting question.

Moving forward, several key avenues for research emerge. First, identifying the neural correlates of geomagnetic (or gravitational) sensitivity in mice and other terrestrial mammals will be critical. Second, understanding how these long-period oscillators interact with circadian and circannual systems could reshape our understanding of the temporal organization of behavior in mammals. Finally, the functional relevance of this semi-synodic modulation remains to be explored: whether it offers adaptive advantages, such as synchronizing reproduction, exploratory, or risk-taking behaviors to predictable environmental cycles. Given the many other parallels seen across humans and rodents, it is possible that infradian rhythms in behavior are a general principle of mammalian biology that has been underexplored due to technological limitations.

## Methods

### Animal husbandry

For this study, we analyzed 9 independent datasets collected under a range of housing and experimental conditions in mice.

#### ColonyRack datasets (D1 - D6)

Six datasets (D1 - D6) were recorded using a custom built modular and automated homecage - system called “ColonyRack” (PhenoSys, Berlin, Germany) ^24^. It consists of 70 individual mouse cages interconnected via polycarbonate tubes, each equipped with radio-frequency identification (RFID) ring antennas at both entry and exit points. Mice were implanted subcutaneously with RFID transponders under brief isoflurane anesthesia prior to the start of experiments. The software PhenoSoft Control logged every antenna contact with a timestamp, animal ID and antenna ID, providing a high - resolution readout of individual mouse movement across the system.

- Datasets D1 and D2 were recorded simultaneously over 166 days from 17 December 2020 to 31 May 2021. Four - week - old female App^NL - G - F/NL - G - F^ (n = 33) and App^NL/NL^ (n = 33) mice were sourced from the RIKEN Institute. These animals carry the Swedish (KM670/671NL), Arctic (E693G), and Beyreuther/Iberian (I716F) mutations associated with Alzheimer’s disease ^70^. After a one - week habituation period, during which RFID transponders were implanted under brief isoflurane anesthesia, the animals were introduced into the ColonyRack system. Mice were maintained under a 12:12 hour light/dark cycle with ad libitum access to food and water at the DZNE animal facility in Dresden. Cages were cleaned weekly. All procedures were conducted in accordance with European and national guidelines and approved by the Landesdirektion Sachsen (TVV 28/2020).
- Datasets D3 and D4 were recorded simultaneously for 86 days from 10 January 2022 to 5 April 2022. The cohort included 4-week old DCX - GFP transgenic mice (40 females, 40 males). Procedures for acclimation, RFID implantation, and housing were identical to those described above. Experiments were conducted at the DZNE Dresden facility under 12:12 hour light/dark conditions with standard environmental parameters and weekly cage cleaning. Ethical approval was granted by the Landesdirektion Sachsen (TVA 16/2018).
- Datasets D5 and D6 were collected simultaneously over 208 days from 7 July 2022 to 31 January 2023. These datasets included female App^NL - F/NL - F^ (n=40) and App^NL/NL^ (n=40) mice carrying the Swedish (KM670/671NL) and Beyreuther/Iberian (I716F) mutations. The same RFID implantation, housing, and maintenance protocols were followed as in D1– D4. Ethical approval was granted by the Landesdirektion Sachsen (TVV 44/2022).

#### Running wheel datasets (R1 - R2)

Datasets R1 and R2 were recorded simultaneously in two different animal facilities in Dresden using a running wheel sensor system (TSE Systems GmbH). Each cage housed either a single mouse or a pair.

- R1 - Female and R1 - Male cohorts were recorded simultaneously for 299 days, from 3 May 2024 to 25 February 2025 at the CRTD animal facility, Dresden. This dataset included C57BL/6JRj mice (female: n = 16; male: n = 15). Mice were maintained on a 12 h light/dark cycle with unrestricted food and water access. Cages were cleaned once per week.
- R2 - Female and R2 - Male cohorts were collected over the same period (3 May 2024 to 5 February 2025, 279 days) at the DZNE animal facility in Dresden. The dataset included C57BL/6JRj mice (female: n = 20; male: n = 13), housed and monitored under identical conditions to the R1 dataset. All animal husbandry and experiments were in accordance with European and national regulations and approved by the local authority (Landesdirektion Sachsen; TVT 3/2020).

#### Data from Acosta-Rodriguez et al., 2022

A previously published dataset from the Takahashi Laboratory (UT Southwestern, Dallas) was included, as described in detail by Acosta-Rodriguez et al., 2022 ^27^. Male C57BL/6J mice, 6 weeks of age, were allowed to acclimatize for 2 weeks before being housed individually in standard cages equipped with stainless steel running wheels. Environmental temperature and humidity were controlled, and animals were maintained under a 12 h light/12 h dark cycle with ad libitum water.

Mice were randomly assigned to one of six feeding conditions. In the ad libitum (AL) group, food was available at all times. In the caloric restriction (CR) groups, 70% of baseline ad libitum intake was provided. This was either given once daily at the beginning of the dark (CR-night-2h) or light (CR-day-2h) phase, which was consumed within approximately 2 h, or distributed evenly over 12 h during the dark (CR-night-12h) or light (CR-day-12h) phase by releasing one 300 mg pellet every 90 min. In the CR-spread condition, 70% of baseline intake was distributed evenly over 24 h by releasing one 300 mg pellet every 160 min.

Every 21 days, the animals were weighed in the morning, cages were changed, and bedding was inspected for food spillage. Feeding schedules were programmed and recorded using ClockLab Chamber Control Software (v3.401, Actimetrics). Wheel-running activity was continuously monitored using the ClockLab Data Acquisition System (v3.209, Actimetrics).

In total, 217 mice were recorded under the different experimental conditions throughout their lifespan, starting from 27 Jan 2017 until their respective dates of death. Across all conditions, there were nine distinct starting dates, resulting in 22 experimental groups. For visualization and analysis, a subset of the data was selected that included only the groups with the starting date 12 Sep 2017, ensuring that all conditions shared a common starting point. The selected dataset covered the period from 12 Sep 2017 to 8 Feb 2021 (1246 days) and included 63 mice (AL: n=5, CR-night-12h: n=12, CR-night-2h: n=11, Cr-day-12h: n=11, CR-day-2h: n=12 and CR-spread: n=12).

#### Data from Izumo et al., 2014

Another previously published dataset from the Takahashi Laboratory (UT Southwestern Medical Center, Dallas) was included, described in detail by Izumo et al. ^28^. The data comprise long-term running wheel activity from eight groups (4 genotypes × 2 sexes) in two cohorts recorded at Evanston, USA from 27 Feb 2007 and 11 Apr 2007 to 24 Sep 2007 (210 days and 166 days). To achieve forebrain - specific disruption of circadian rhythms, *Bmal1*^fx/fx^ mice ^71^ were crossed with *Camk2a::iCre* BAC mice^72^, generating *CamiCre+; Bmal1*^fx/fx^ (cKO; female, n=8; male, n=10) mice and three control genotypes: *Bmal1*^fx/fx^ (FxFx; female, n=9; male, n=9), *CamiCre*+ (Cre; female, n=8; male, n=5), and *CamiCre+; Bmal1*^fx/+^ (Het; female, n=8; male, n=8). At >8 weeks of age, mice were individually housed in cages equipped with activity wheels and monitored under a 12 h light/dark (LD) cycle for at least 10 days. Subsequently, animals were transferred to constant darkness (DD) for 4 weeks, returned to LD for 2 weeks, exposed to constant light (LL) for 4 weeks, and finally re-introduced to LD for the rest of the experiment. Locomotor activity was recorded using ClockLab software (Actimetrics, Wilmette, IL).

### Behavioral analysis

For the ColonyRack datasets (D1 - D6), PhenoSoft Control software outputs a raw data file containing tabular information about each animal’s visit to individual cages through antenna contacts. The exploratory activity was examined by calculating roaming entropy as described previously ^25,26^. Briefly, for each mouse, the amount of time spent in each cage was calculated by summing up the durations between successive antenna contacts. These durations were assigned to cages based on the antenna’s physical location. From this, the proportion of time spent in each cage (cage probability) was calculated as *p_i_*_,*j*,*t*_ of a mouse *i* being in a cage *j* in a time bin *t*. Using these probabilities, Shannon entropy of the roaming distribution was calculated as

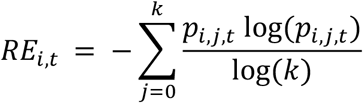

where k is the number of total cages. Normalizing the entropy by log(k) scales the RE between 0 (indicating absolute spatial bias, time spent in only one cage) and 1 (indicating a perfectly uniform distribution of time across all cages). The mean RE was subsequently computed across all animals within each dataset, providing a group - level measure.

For running wheel datasets (R1, R2 and data from Acosta-Rodriguez et al., 2022 and Izumo et al., 2014), locomotor activity was quantified by calculating the total number of wheel rotations recorded per cage.

### Time series analysis

#### Data cleaning

Missing values due to technical interruptions in the mean RE and mean activity time series were imputed using a seasonal decomposition - based interpolation method. A moving average algorithm was applied to estimate and replace the missing values, ensuring that the underlying temporal structure of the data was preserved. This approach was implemented using the imputeTS package in R ^73^. A linear detrending procedure was then applied to the time series by first identifying a linear trend estimated via regression on time and subtracted from the original mean time series.

#### Spectral analysis

To examine the underlying periodic structure in the detrended mean time series, a spectral density analysis was performed using the ‘spectrum’ package in R ^74^. Finally, for plotting periodograms, spectral density values were min - max normalized to a 0 - 1 range for consistent comparison across datasets. Peaks in the periodograms were detected using the ‘findpeaks’ function from ‘pracma’ package in R ^75^.

Using wavelet transforms, time-varying periodicities in the signal were examined using ‘WaveletComp’ R package ^76^. The input time series was first smoothed with a loess regression to reduce noise. The resulting wavelet power spectrogram was visualized as a time - period heatmap, illustrating how the signal’s dominant periodic components evolve over time. Finally, peaks in the average wavelet power spectrum were detected to identify the principal dominant periods present in the data.

#### Harmonic reconstruction of waveforms from frequency - domain components

A custom function was used to reconstruct time-domain waveforms from frequency-domain data for visualization and peak detection. Peaks with periods exceeding 10 days were first identified within the frequency spectrum, and their corresponding amplitudes and phases were extracted. Following the inverse Fourier approach, at each time point, the waveform was generated by summing sinusoidal components weighted by their respective amplitudes and phase offsets.

#### Autocorrelation analysis

To further examine the periodic structure in the behavioral metrics, an autocorrelation analysis was performed on the detrended mean time series at a 24 - hour resolution. The autocorrelation was computed using the ‘ccf’ function from ‘tseries’ package in R ^77^, computing correlations across lag values up to the length of the time series minus one, with a lag interval of one day. The resulting autocorrelation coefficients were plotted across lags to visualize the strength and periodicity in the behavioral fluctuations.

### Analysis of lunar modulation of behavioral metrics

#### Extraction of lunar data

Lunar metrics corresponding to each date in the datasets were extracted using functions from the ‘lunar’ ^78^ and ‘suncalc’ ^79^ packages in R. The extracted parameters included the fractional illumination of the Moon (0 to1), the synodic phase expressed as a continuous measure in radians, and 8 - category classification of the lunar phase (New Moon, Waxing Crescent, First Quarter, Waxing Gibbous, Full Moon, Waning Gibbous and Last Quarter). Additionally, the Earth–Moon distance was calculated for each date, expressed as a continuous value in Earth radii indicating proximity to perigee (∼56 Earth radii) or apogee (∼63 Earth radii). To calculate the phase of the anomalistic cycle i.e., Moon’s orbit relative to its distance from Earth, peaks and valleys in the Earth–Moon distance timeseries were identified. Peaks correspond to apogee (the Moon’s farthest point from Earth), and valleys to perigee (the closest point). These extrema were marked with angular phase values: π at apogee and 2π at perigee. Intermediate phase values between these reference points were linearly interpolated, resulting in a continuous measure of the Moon’s anomalistic phase expressed in radians over time. These variables were compiled into a single dataset for subsequent examination of potential lunar modulation in behavioral patterns.

#### Quantifying lunar - phase effects on behavior

To assess phase-dependent modulation of behavior across the lunar synodic cycle, the continuous synodic phase (0 to 2π radians) was divided into 28 equal-width bins. Each time point was assigned to a bin based on its phase value. For each bin, mean values of behavioral metrics (RE and activity) and lunar distance were computed, followed by min - max scaling from 0–1. Corresponding angular positions for each bin were calculated to create graphical representation with bin aggregated behavioral activity and lunar distance plotted against synodic phase. For visualization, the x - axis was scaled to cover two complete synodic cycles (0 to 4π radians).

#### Cross - Wavelet Power analysis

The ‘WaveletComp’ R package ^76^ was used to compute cross-wavelet power between behavior and lunar time series. This method quantifies the shared spectral power across time and frequency domains, identifying periods of strong joint oscillatory activity.

#### Quantification of behavioral peak alignments with the lunar cycle

To quantify the alignment of behavioral peaks with lunar events, peak times from the reconstructed waveforms were compared to known full and new moon dates. Matched time points were first used to calculate the difference in days between behavioral peak and the closest full or new moon event (Δ*Days*). These matched time points were then converted into circular phase values normalised on the lunar period (29.53 days) and expressed in radians.

The Phase Locking Value (PLV) was then calculated as the magnitude of the mean resultant vector of these phase differences:

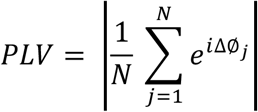

where *N* is the number of peak-moon event pairs and ΔØ_*j*_ is the phase difference. The PLV ranges from 0 (no phase alignment) to 1 (perfect alignment).

The mean phase angle was computed as the argument of the mean vector

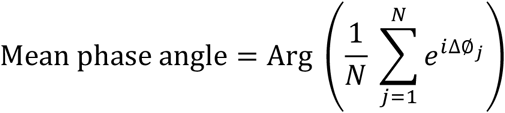

representing the average timing of peaks relative to the lunar reference. If the mean phase angle equals 0, behavioral peaks occur on average exactly at the reference event; positive values indicate peaks occurring after the reference, while negative values indicate peaks occurring before it.

To test whether this observed synchrony could have happened by chance, a permutation test was run. Over 1,000 iterations, the peak times from one signal were randomly shuffled to break any temporal alignment. For each shuffled set, phase differences and the corresponding PLV were recalculated. The final p - value was determined by checking how many of these permuted PLVs were equal to or greater than the observed PLV, giving an estimate of the likelihood that the observed alignment could occur by chance.

#### Rayleigh test for unimodal and bimodal phase locking

To assess the alignment of behavioral peaks with specific phases of the synodic lunar cycle, circular statistical analyses were conducted separately for each dataset using the ‘circular’ package in R ^80^. First, smoothed trajectories of behavioral metrics were generated using harmonic reconstruction of the time series from frequency - domain components. Peaks were then identified from these smoothed waveforms, and their corresponding phase values (in radians) were determined based on their position within the lunar cycle. These values were treated as angular data and converted into circular format.

To evaluate whether the phase distribution deviated from uniformity (i.e., whether behavioral peaks preferentially occurred around a specific lunar phase), the standard Rayleigh test for unimodal clustering was applied. Since visual inspection of the data suggested that peaks tended to cluster around both the full and new moon indicative of axial bimodality, a second Rayleigh test was performed after doubling the phase angles. This transformation effectively converts an axial bimodal distribution into a unimodal one, allowing the Rayleigh test to assess bimodal clustering as well.

### Extraction of observed geomagnetic variations

Geomagnetic field data from ground geophysical observatories were retrieved from publicly available databases. Specifically, we used the *VirES for Swarm* Python software ^81^ that delivers (quasi-)definitive data from INTERMAGNET and the World Data Centre (WDC) for Geomagnetism, following the quality control method by ^82^. Hourly means of the vector geomagnetic field, measured at the nearest geomagnetic observatory to the location of the behavioral experiments, were used for the experimental periods.

The total geomagnetic field contains contributions from various internal and external magnetic field sources, which vary at different, albeit overlapping, spatial and temporal scales ^83^. The total field intensity is dominated by the core field that accounts for up to 95-99% of its magnitude. However, the core field varies slowly over time (from annual to secular scales). The other field sources are magnetized rocks in the Earth’s crust. This component is virtually constant, varying at millennial scales. To focus on geomagnetic field variations that originate in the magnetosphere and ionosphere (collectively referred to as external field sources) and vary at periods of interest, we therefore subtracted the core and crustal field components as given by the CHAOS-8 model ^84^ from the observed field.

Although all vector components of the geomagnetic field are sensitive to ionospheric and magnetospheric variations, the magnitude of the external geomagnetic variations is often most pronounced in the North component of the geomagnetic field. Therefore, only this component was used in the geomagnetic data analysis.

The hourly geomagnetic time series was used for spectral analyses to identify dominant periodic components. To examine possible coupling between geomagnetic variations and behavioral fluctuations, daily means for both the time series were used to calculate cross-correlation and phase-synchronization measures (PLV).

## Data availability

The ColonyRack datasets (D1-D4) recorded in Dresden, together with the corresponding geomagnetic-field data recorded at the Niemegk geomagnetic observatory, are available on figshare with the identifier: 10.6084/m9.figshare.32592147 (The figshare repository will be made publicly available before publication; in the meantime, it can be accessed via the following private link: https://figshare.com/s/caf33655c462def45619). Other cohorts and external datasets described in the manuscript are not included in this figshare record; access to those datasets may be requested from the authors of the respective original studies.

## Code availability

The R scripts used to process the ColonyRack behavioral data, analyze lunar and geomagnetic alignment, and reproduce the corresponding figures are available on figshare with the identifier: 10.6084/m9.figshare.32592147.

## Acknowledgments

W.B. was supported by the International Max Planck Research School on Learning, Institutions, and Future Evolution (LIFE, www.imprs-life.mpg.de; participating institutions: Max Planck Institute for Human Development, Freie Universität Berlin, Humboldt-Universität zu Berlin, Technische Universität Berlin, University of Michigan, University of Virginia, University of Zurich) and by the Federal Ministry of Research, Technology and Space (BMFTR) as part of the German Center for Child and Adolescent Health (DZKJ; funding code 01GL2405B). We thank members of the Kempermann lab for assistance with the animal experiments and care; S. Guenther, A. Karasinsky and J. Bergmann. We are grateful to F. Ehret and H. Liu for sharing experimental data from the App^NL-G-F^ and App^NL-F^ experiments respectively. V.A.R was supported by the Intramural Research Program of the National Institutes of Health (NIH). The contributions of the NIH author(s) were made as part of their official duties as NIH federal employees, are in compliance with agency policy requirements, and are considered Works of the United States Government. However, the findings and conclusions presented in this paper are those of the author(s) and do not necessarily reflect the views of the NIH or the U.S. Department of Health and Human Services. A.G. was supported by the Heisenberg Grant from the German Research Foundation, Deutsche Forschungsgemeinschaft (project no. 465486300), and the ESA Swarm DISC project no. 4000109587. J.S.T. was an Investigator in the Howard Hughes Medical Institute when experiments were conducted. G.K. was supported by the grant “The mouse in the supermarket” from Volkswagen Foundation.

## Author contributions

Conceptualization: WB, AER, GK

Methodology: WB, AG, MI, VAR, JST, GK

Visualization and analysis: WB, AG

Funding acquisition: JST, GK

Project administration: AER, GK

Supervision: AER, JST, GK

Writing – original draft: WB, GK

Writing – review & editing: WB, AG, AER, MI, VAR, JST, GK

## Supplementary Materials

Figs. S1 to S19

Tables S1 to S5

