## Supplemental Material for "Mice sense Moon and Sun"

### Mice sense the Sun and the Moon

**The PDF file includes:**

Figs. S1 to S9
Tables S1 to S5

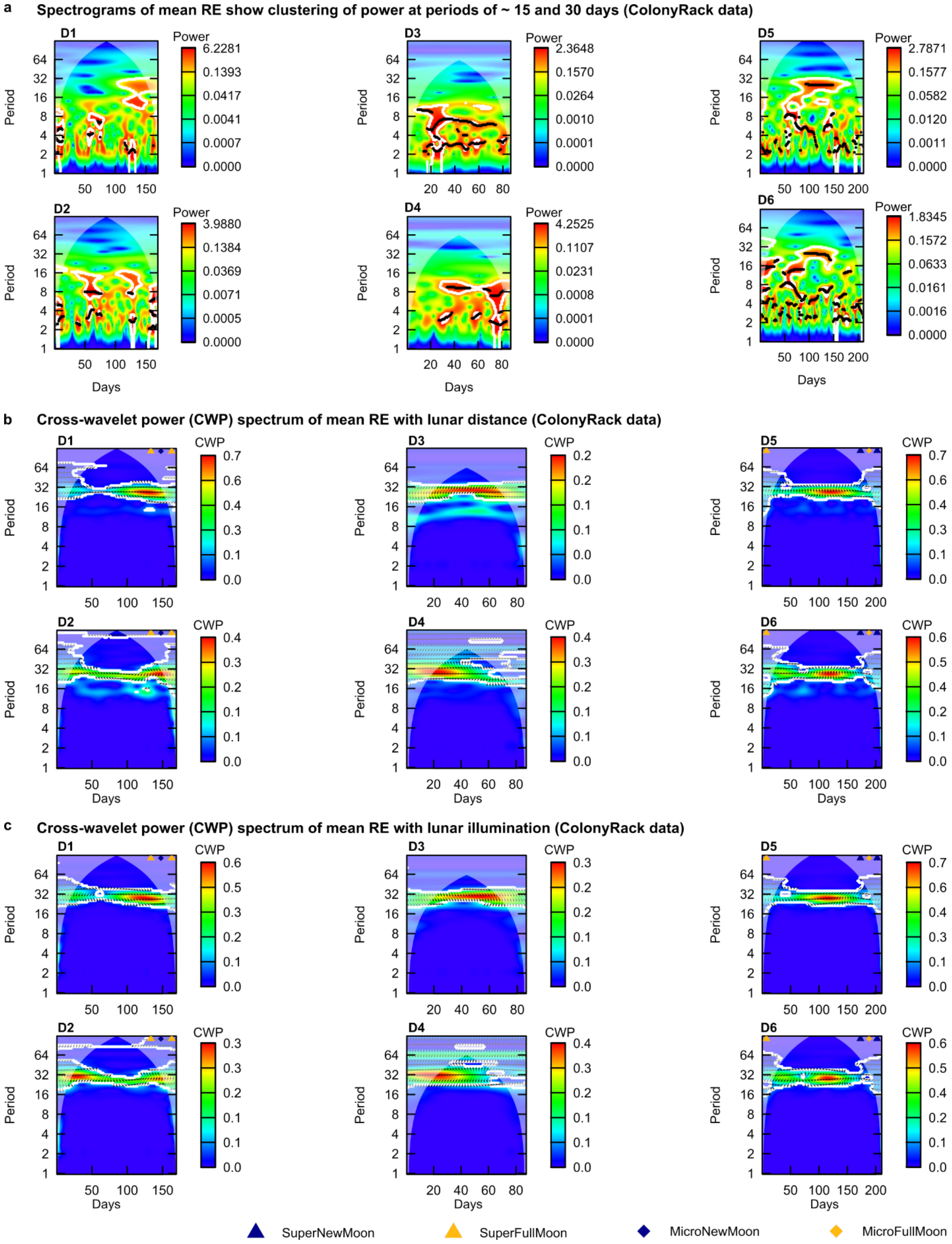
a Spectrograms of mean RE show clustering of power at periods of ~ 15 and 30 days (ColonyRack data)
b Cross-wavelet power (CWP) spectrum of mean RE with lunar distance (ColonyRack data)
c Cross-wavelet power (CWP) spectrum of mean RE with lunar illumination (ColonyRack data)
D1
D2
D3
D4
D5
D6
Period
Days
Power
CWP
SuperNewMoon
SuperFullMoon
MicroNewMoon
MicroFullMoon

**Fig. S1. Spectral analysis of mean RE across ColonyRack datasets D1 – D6.**

**(a)** Spectrograms of mean RE. Peak positions are reported in Table S1. **(b)** Cross - wavelet power spectrogram of mean RE with lunar distance. For information about peak positions see Table S2. **(c)** Cross-wavelet power spectrogram of mean RE with lunar illumination. For information about peak positions see Table S2.

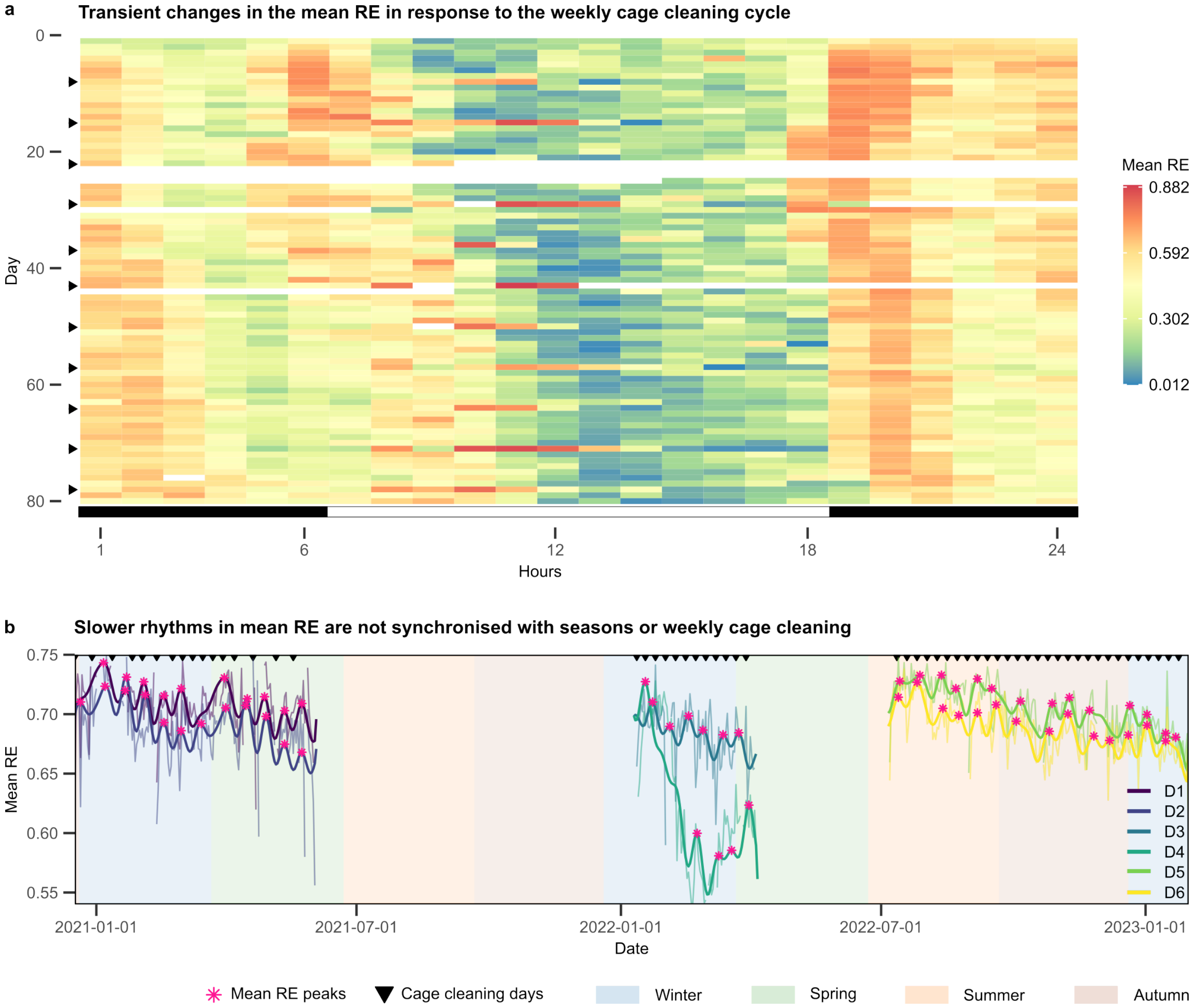


**Fig. S2. Infradian rhythms in mean RE are not modulated by weekly cage cleaning cycles or seasonal changes.**

**(a)** Actogram showing mean RE across 40 animals from the ColonyRack dataset D3. Actogram shows rows corresponding to the days of the experiment and columns for 24 hours of the day, color-coded to indicate RE levels. Missing values are colored as white. Weekly cage changes (black arrows on the left) trigger brief increases in activity that subside within a few hours. **(b)** Datasets D1-D6 span different seasons over multiple years. Mean RE does not show modulation by seasons or seasonal transitions. Raw waveforms (plotted with transparent colors) and reconstructed slow waveforms (bold lines) are shown against a background indicating the seasons of the year. Local peaks of the slow waveform are marked with asterisks. The continuous bold line for each dataset represents a reconstructed waveform, generated by summing sine waves based on the dominant frequencies, amplitudes, and phases identified in the Fourier spectrum. Weekly cage cleaning days, which elicit a transient weekly response in the raw waveform, are indicated by inverted black triangles.

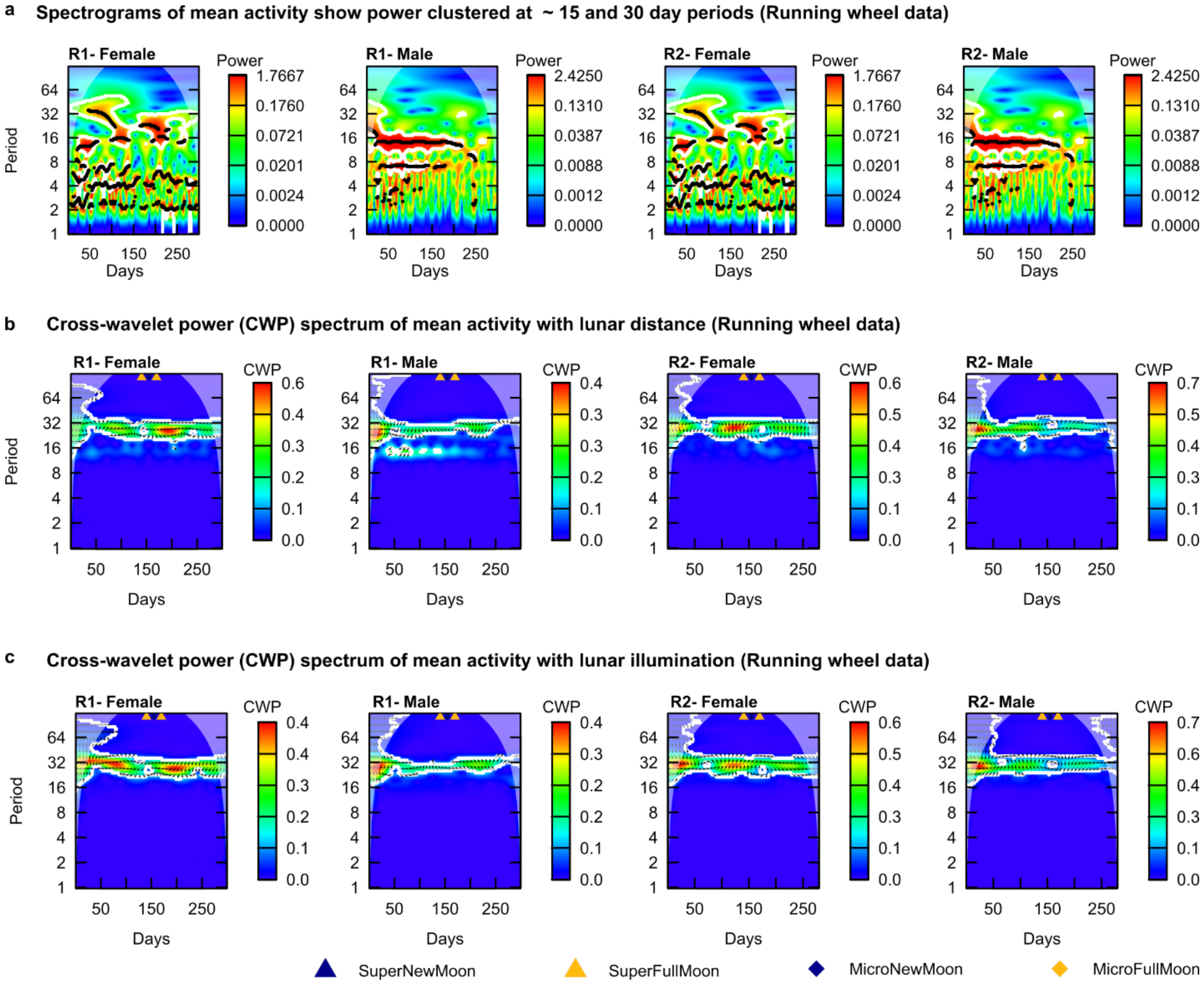


**Fig. S3. Spectral analysis of mean activity across running wheel datasets R1 – R2.**

**(a)** Spectrograms of mean activity. Peak positions are reported in Table S1. **(b)** Cross - wavelet power spectrogram of mean activity with lunar distance. For information about peak positions see Table S2. **(c)** Cross-wavelet power spectrogram of mean activity with lunar illumination. For information about peak positions see Table S2.

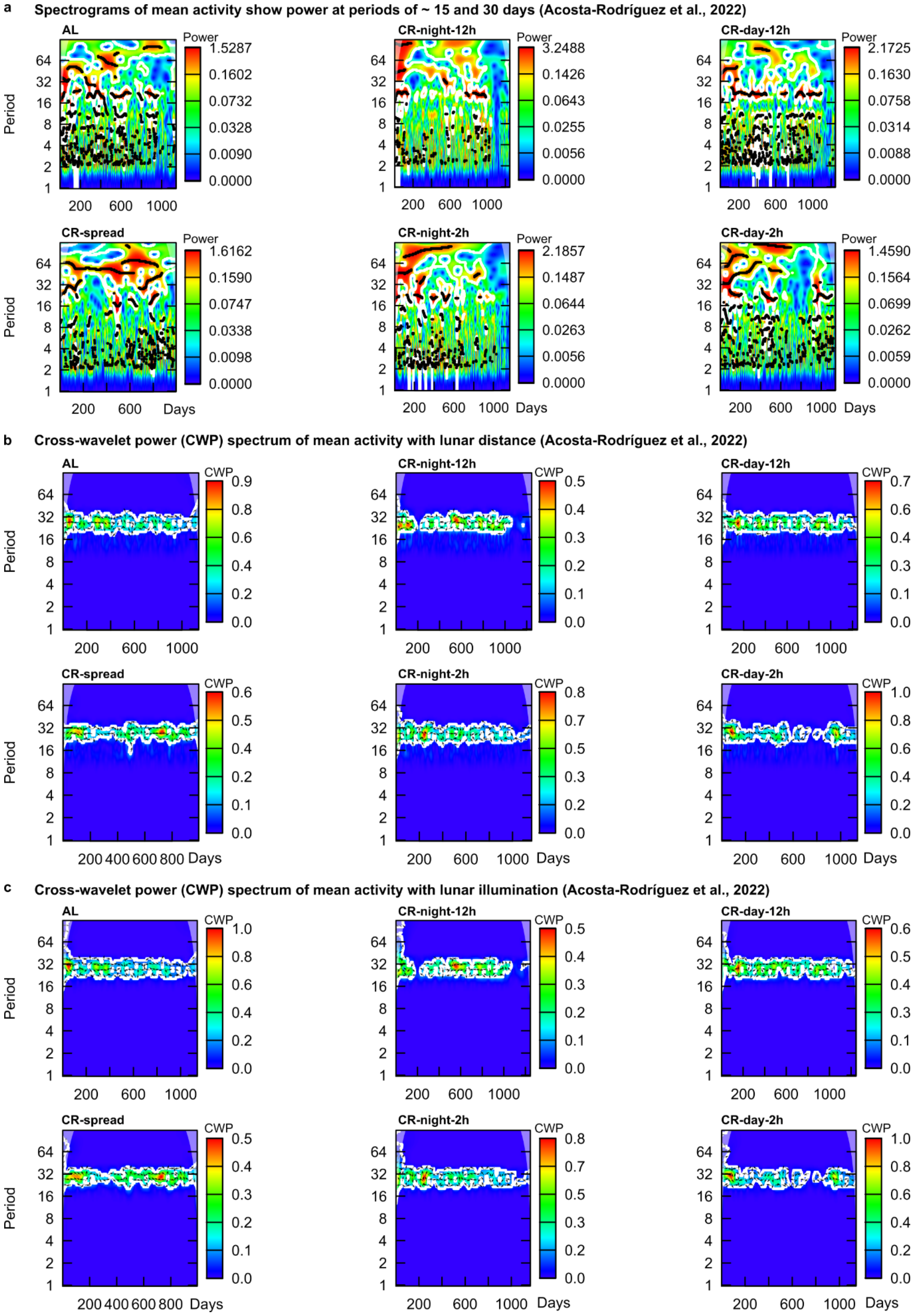
a
Spectrograms of mean activity show power at periods of ~ 15 and 30 days (Acosta-Rodríguez et al., 2022)
AL
CR-night-12h
CR-day-12h
CR-spread
CR-night-2h
CR-day-2h
Power
Period
Days
b
Cross-wavelet power (CWP) spectrum of mean activity with lunar distance (Acosta-Rodríguez et al., 2022)
AL
CR-night-12h
CR-day-12h
CR-spread
CR-night-2h
CR-day-2h
CWP
Period
Days
c
Cross-wavelet power (CWP) spectrum of mean activity with lunar illumination (Acosta-Rodríguez et al., 2022)
AL
CR-night-12h
CR-day-12h
CR-spread
CR-night-2h
CR-day-2h
CWP
Period
Days

**Fig. S4. Spectral analysis of mean activity across running wheel datasets from Acosta-Rodriguez et al., 2022.**

**(a)** Spectrograms of mean activity. Peak positions are reported in Table S1. **(b)** Cross - wavelet power spectrogram of mean activity with lunar distance. For information about peak positions see Table S2. **(c)** Cross - wavelet power spectrogram of mean activity with lunar illumination. For information about peak positions see Table S2.

**a** **Spectrograms of mean activity show power at periods of ~ 15 and 30 days (Izumo et al., 2014 data)**

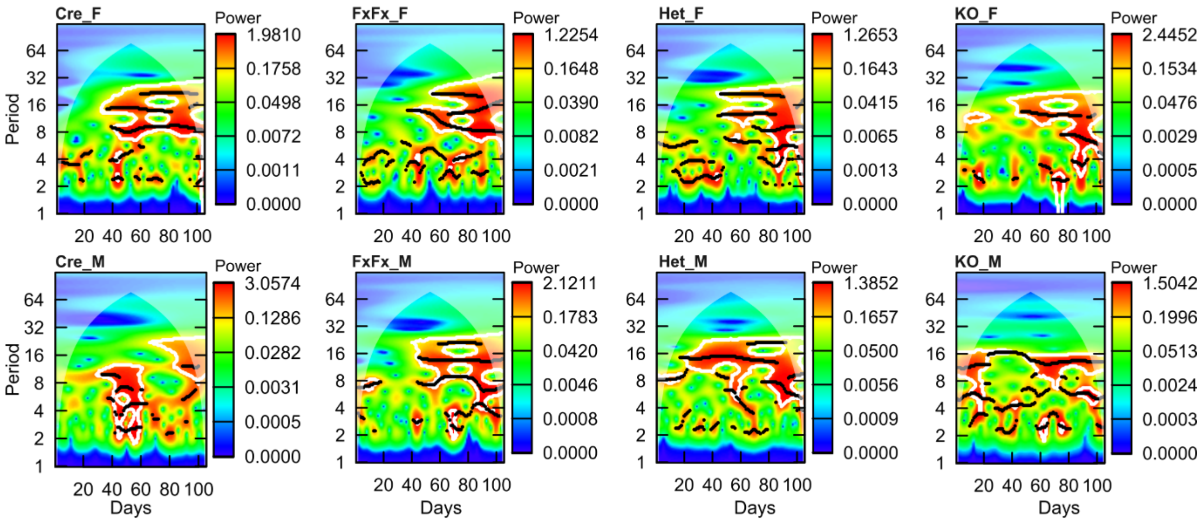


**b** **Cross-wavelet power (CWP) spectrum of mean activity with lunar distance (Izumo et al., 2014 data)**

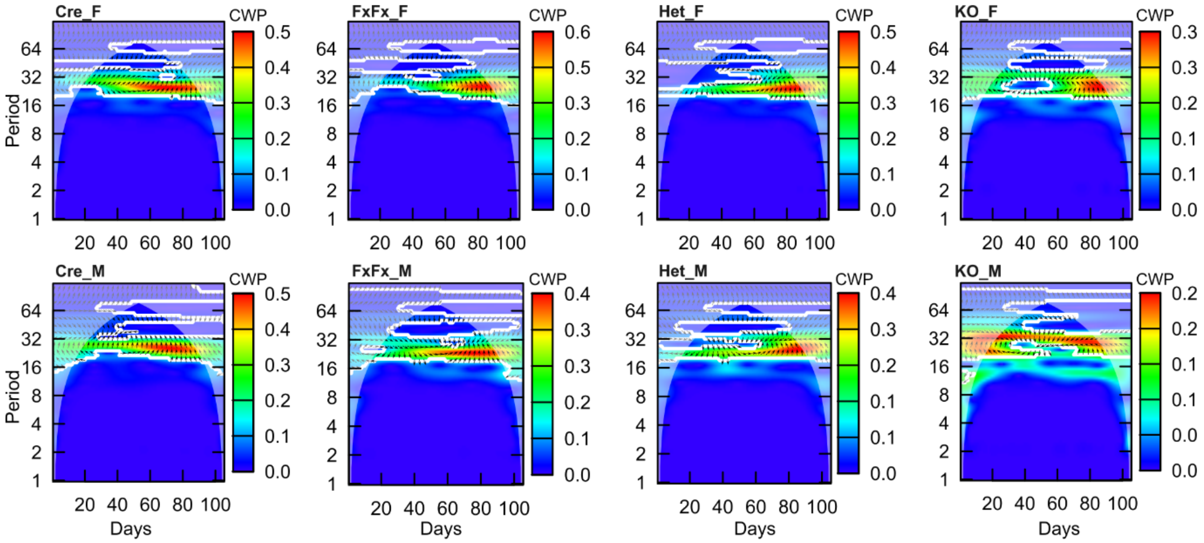


**c** **Cross-wavelet power (CWP) spectrum of mean activity with lunar illumination (Izumo et al., 2014 data)**

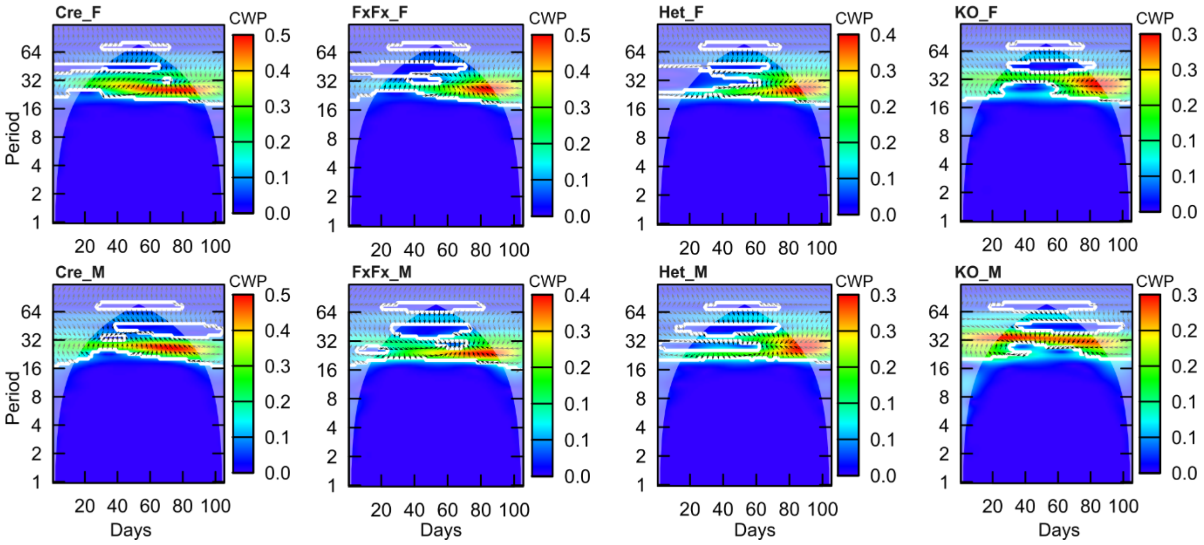

**Fig. S5. Spectral analysis of mean activity across running wheel datasets from Izumo et al., 2014.**

**(a)** Spectrograms of mean activity. Peak positions are reported in Table S1. **(b)** Cross - wavelet power spectrogram of mean activity with lunar distance. For information about peak positions see Table S2. **(c)** Cross - wavelet power spectrogram of mean activity with lunar illumination. For information about peak positions see Table S2.

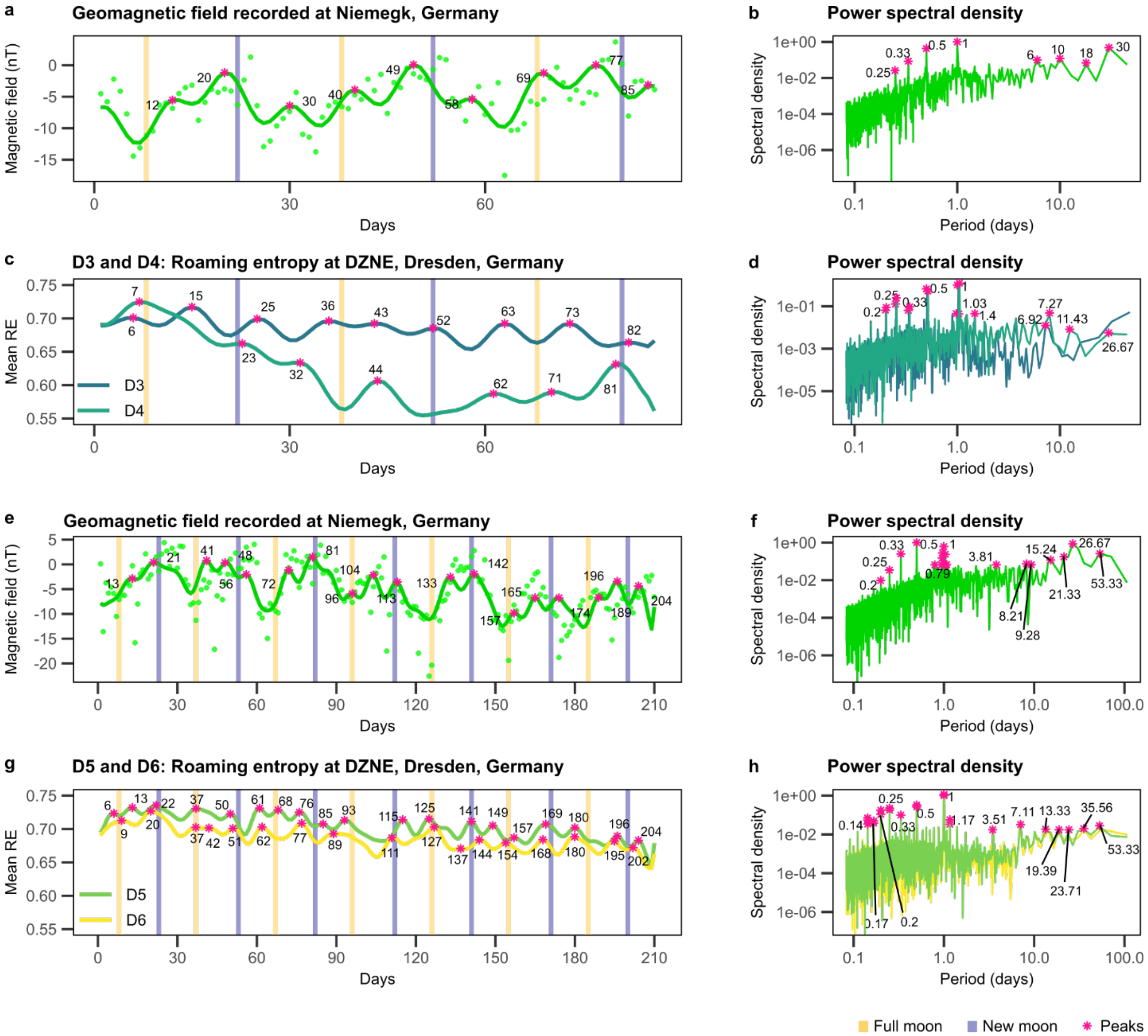


**Fig. S6. D3-D6: Mean RE and local geomagnetic field intensity co-vary with overlapping dominant frequencies.**

**(a, b)** Geomagnetic field recorded at Niemegk observatory, Germany from 10 Jan to 5 Apr 2022 (86 days). **(c, d)** Datasets D3 and D4: RE in DCX-GFP mice (40 females, 40 males) recorded concurrently, with corresponding power spectral density. **(e, f)** Geomagnetic field recorded at Niemegk observatory, Germany from 7 Jul 2022 to 31 Jan 2023 (208 days). **(g, h)** Datasets D5 and D6: RE in App$^{NL-F/NL-F}$ (n = 40) and App$^{NL/NL}$ (n = 40) mice over the same period, with corresponding power spectral density. Across datasets, behavioral and geomagnetic signals exhibit temporally aligned peaks, with shared circadian and infradian spectral components. Jitter in alignment, phase locking values, and corresponding statistics are reported in Table S5.

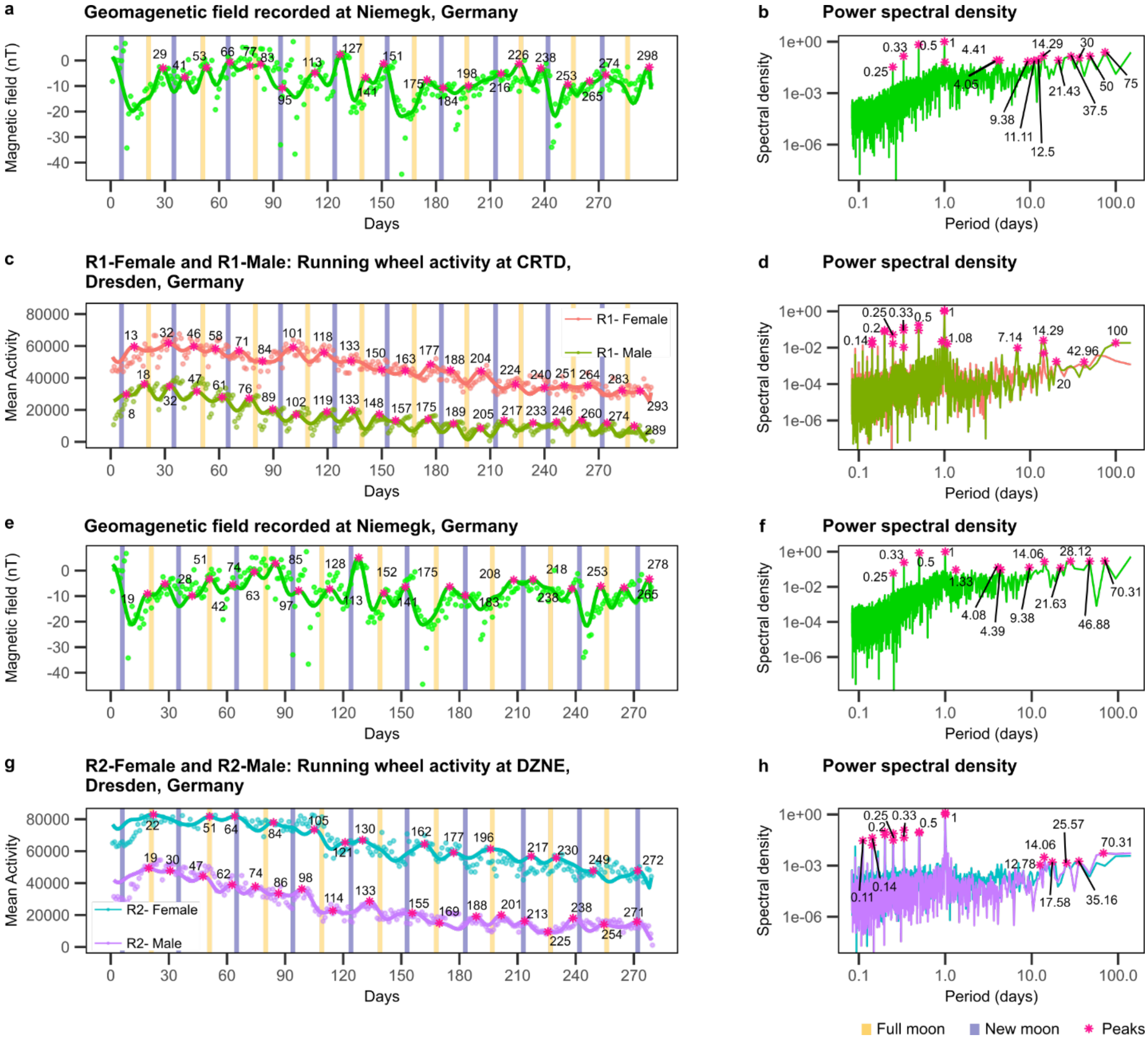

**Fig. S7. R1-R2: Mean activity and local geomagnetic field intensity co-vary with overlapping dominant frequencies.**

**(a, b)** Geomagnetic field recorded at Niemegk observatory, Germany, from 3 May 2024 to 25 February 2025 (299 days), with corresponding power spectral density. **(c, d)** Dataset R1-Female and R1-Male: Mean activity in C57BL/6JRj mice (female: n = 16; male: n = 15) recorded at CTRD, Dresden, over the same period, with corresponding power spectral density. **(e, f)** Geomagnetic field from 3 May 2024 to 5 February 2025 (279 days). **(g, h)** Dataset R2-Female and R2-Male: Mean activity in C57BL/6JRj mice (female: n = 20; male: n = 13) recorded concurrently at DZNE, Dresden, with corresponding power spectral density. Across datasets, behavioral and geomagnetic signals exhibit temporally aligned peaks, with shared circadian and infradian spectral components. Jitter in alignment, phase locking values, and corresponding statistics are reported in Table S5.

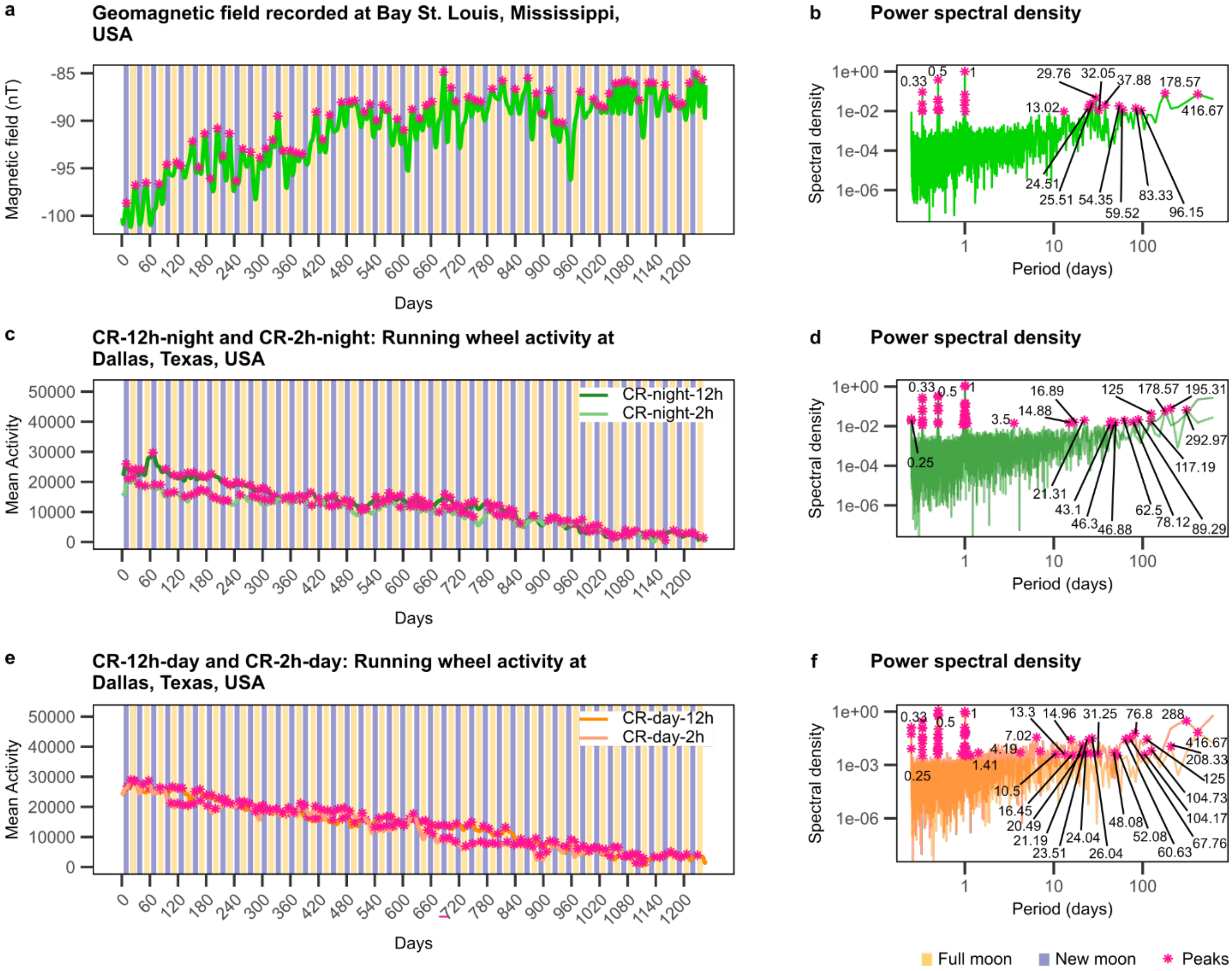


**Fig. S8. Acosta-Rodriguez et al., 2022: Mean activity and local geomagnetic field intensity co-vary with overlapping dominant frequencies.**

**(a, b)** Geomagnetic field recorded at Bay St. Louis observatory, USA, from 12 Sep 2017 to 8 Feb 2021 (1246 days) with corresponding power spectral density. **(c, d)** Dataset CR-night-12h and CR-night-2h: Mean activity in C57BL/6J mice (CR-night-12h: n = 12; CR-night-2h: n = 11) recorded at Dallas, USA over the same period, with corresponding power spectral density. **(e, f)** Dataset CR-day-12h and CR-day-2h: Mean activity in C57BL/6J mice (CR-day-12h: n = 11; CR-day-2h: n = 12) recorded at Dallas, USA over the same period, with corresponding power spectral density. Across datasets, behavioral and geomagnetic signals exhibit temporally aligned peaks, with shared circadian and infradian spectral components. Jitter in alignment, phase locking values, and corresponding statistics are reported in Table S5.

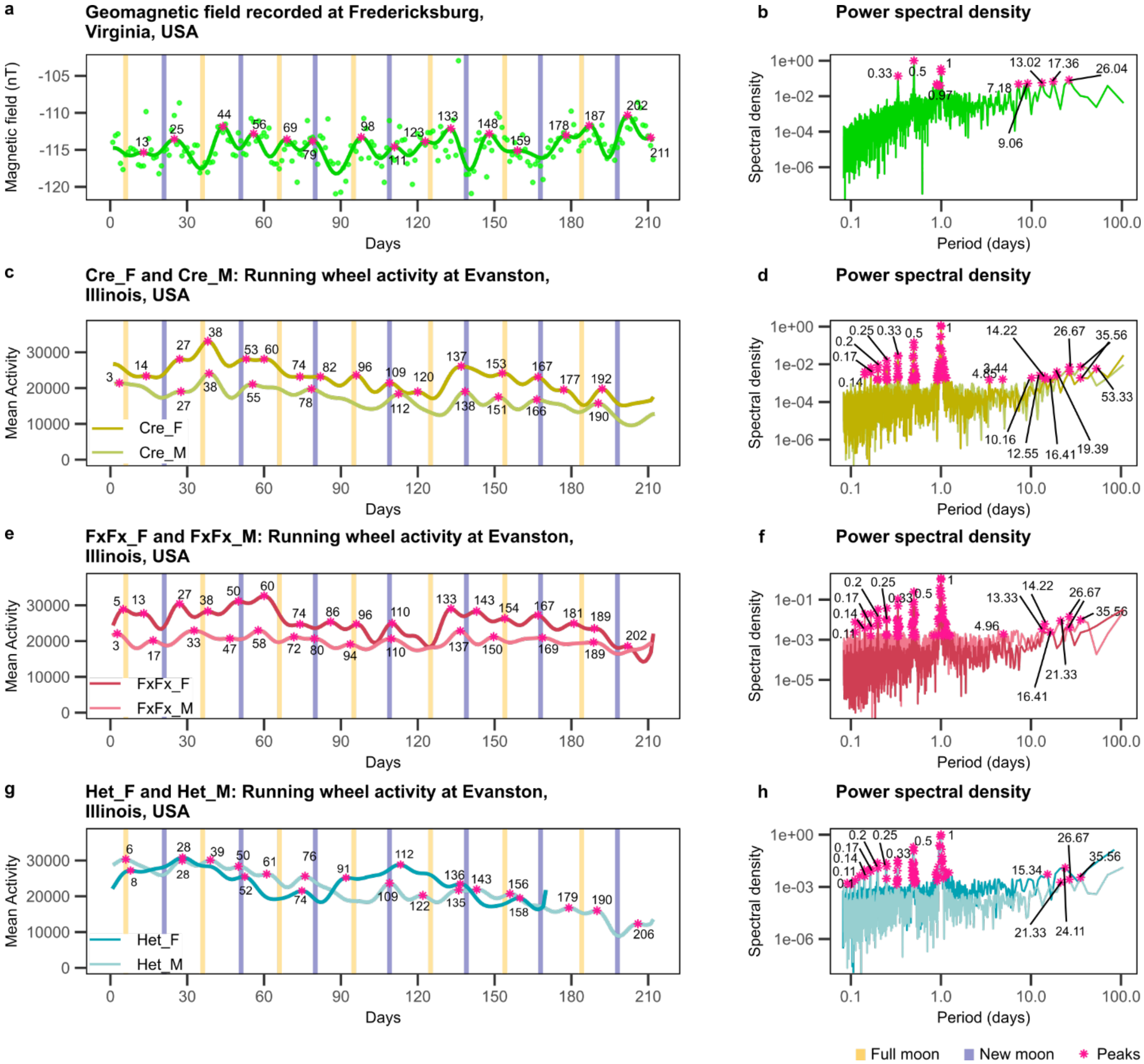


**Fig. S9. Izumo et al., 2014: Mean activity and local geomagnetic field intensity co-vary with overlapping dominant frequencies.**

**(a, b)** Geomagnetic field recorded at Fredricksburg observatory, USA, from 27 Feb 2007 and 11 Apr 2007 to 24 Sep 2007 (210 days), with corresponding power spectral density. **(c, d)** Dataset Cre: Mean activity in *CamiCre*+ mice (Cre; female n=8; male, n=5) recorded at Evanston, USA over the same period, with corresponding power spectral density. **(e, f)** Dataset FxFx: Mean activity in *Bmal1*fx/fx mice (FxFx; female, n=9; male, n=9) recorded at Evanston, USA over the same period, with corresponding power spectral density. **(g, h)** Dataset Het: Mean activity in *CamiCre*+; *Bmal1*fx/+ mice (Het; female, n=8; male, n=8) recorded at Evanston, USA over the same period, with corresponding power spectral density. Across datasets, behavioral and geomagnetic signals exhibit temporally aligned peaks, with shared circadian and infradian spectral components. Jitter in alignment, phase locking values, and corresponding statistics are reported in Table S5.

**Table S1: Position of peaks in mean RE/activity spectrograms**

| Data | 1st Dominant period (days) | 2nd Dominant period (days) |
|---|---|---|
| **D1** | 25.10 | 13.50 |
| **D2** | 26.90 | 13.90 |
| **D3** | 32.00 | 10.20 |
| **D4** | 18.40 | 18.40 |
| **D5** | 25.10 | 13.90 |
| **D6** | 26.00 | 13.00 |
| **R1 - F** | 28.84 | 14.42 |
| **R1 - M** | 30.91 | 13.93 |
| **R2 - F** | 29.86 | 16.56 |
| **R2 - M** | 26.91 | 15.45 |
| **Cre - F** | 21.11 | 14.42 |
| **Cre - M** | 20.39 | 9.85 |
| **FxFx - F** | 20.39 | 13.45 |
| **FxFx - M** | 20.39 | 13.45 |
| **Het - F** | 20.39 | 12.55 |
| **Het - M** | 13.93 | 13.93 |
| **cKO - F** | 18.38 | 12.13 |
| **cKO - M** | 30.91 | 13.00 |
| **AL** | 26.90 | 13.50 |
| **CR-12h-day** | 19.70 | 13.50 |
| **CR-12h-night** | 21.11 | 13.45 |
| **CR-2h-day** | 22.63 | 14.42 |
| **CR-2h-night** | 32.00 | 19.70 |
| **CR-spread** | 27.86 | 13.45 |

**Table S2: Position of peaks in cross wavelet transforms between mean RE/activity with lunar distance and lunar illumination.**

| | Lunar Distance | | Lunar Illumination | |
|---|---|---|---|---|
| **Data** | **1st Dominant period (days)** | **2nd Dominant period (days)** | **1st Dominant period (days)** | **2nd Dominant period (days)** |
| **D1** | 26.90 | 14.30 | 28.50 | 8.98 |
| **D2** | 26.90 | 14.30 | 28.50 | 5.99 |
| **D3** | 26.90 | 12.70 | 28.50 | 11.99 |
| **D4** | 28.50 | 12.70 | 32.00 | 4.24 |
| **D5** | 26.90 | 14.30 | 26.90 | 6.73 |
| **D6** | 26.90 | 14.30 | 26.90 | 4.49 |
| **R1 - F** | 28.51 | 14.25 | 30.20 | 4.49 |
| **R1 - M** | 28.51 | 14.25 | 30.20 | 5.34 |
| **R2 - F** | 28.51 | 16.00 | 30.20 | 7.55 |
| **R2 - M** | 26.91 | 15.10 | 28.51 | 5.99 |
| **Cre - F** | 25.40 | 14.25 | 26.91 | 9.51 |
| **Cre - M** | 26.91 | 7.13 | 26.91 | 7.13 |
| **FxFx - F** | 26.91 | 14.25 | 26.91 | 10.08 |
| **FxFx - M** | 22.63 | 14.25 | 23.97 | 7.55 |
| **Het - F** | 23.97 | 14.25 | 23.97 | 10.08 |
| **Het - M** | 23.97 | 14.25 | 23.97 | 10.08 |
| **cKO - F** | 22.63 | 13.45 | 23.97 | 5.99 |
| **cKO - M** | 22.63 | 14.25 | 22.63 | 5.66 |
| **AL** | 26.91 | 15.10 | 30.20 | 5.99 |
| **CR-12h-day** | 25.40 | 3.56 | 30.20 | 3.56 |
| **CR-12h-night** | 23.97 | 15.10 | 30.20 | 4.49 |
| **CR-2h-day** | 26.91 | 15.10 | 28.51 | 2.24 |
| **CR-2h-night** | 25.40 | 15.10 | 30.20 | 7.55 |
| **CR-spread** | 26.91 | 15.10 | 28.51 | 6.73 |

**Table S3: Rayleigh test for mean RE and activity.**

| | **Synodic phase** | | **Anomalistic phase** | |
|---|---|---|---|---|
| **Data** | **Unimodal p-value** | **Bimodal p-value** | **Unimodal p-value** | **Bimodal p-value** |
| **D1** | 2.06E - 01 | 3.05E - 10 | 4.34E - 01 | 7.99E - 03 |
| **D2** | 6.24E - 01 | 1.78E - 09 | 8.35E - 01 | 7.49E - 03 |
| **D3** | 8.33E - 02 | 1.15E - 01 | 2.93E - 01 | 6.93E - 01 |
| **D4** | 3.63E - 01 | 5.57E - 01 | 4.03E - 01 | 6.70E - 01 |
| **D5** | 2.55E - 02 | 4.41E - 04 | 9.67E - 01 | 7.12E - 03 |
| **D6** | 3.66E - 01 | 1.09E - 02 | 9.72E - 01 | 1.58E - 03 |
| **R1 - F** | 2.67E - 02 | 7.08E - 04 | 4.94E - 01 | 8.06E - 01 |
| **R1 - M** | 8.63E - 02 | 3.14E - 05 | 8.35E - 01 | 2.01E - 01 |
| **R2 - F** | 7.99E - 01 | 1.81E - 01 | 9.92E - 01 | 5.91E - 10 |
| **R2 - M** | 7.68E - 01 | 3.04E - 03 | 6.65E - 01 | 3.33E - 02 |
| **Cre - F** | 7.54E - 01 | 3.50E - 05 | 5.47E - 01 | 8.46E - 07 |
| **Cre - M** | 6.07E - 03 | 9.03E - 02 | 8.60E - 02 | 9.97E - 01 |
| **FxFx - F** | 2.50E - 03 | 5.26E - 02 | 6.17E - 04 | 4.15E - 01 |
| **FxFx - M** | 8.40E - 01 | 3.64E - 05 | 4.99E - 01 | 4.16E - 04 |
| **Het - F** | 8.84E - 01 | 2.22E - 02 | 6.37E - 01 | 3.40E - 02 |
| **Het - M** | 5.46E - 01 | 6.10E - 02 | 4.35E - 02 | 3.95E - 02 |
| **cKO - F** | 6.32E - 01 | 4.72E - 02 | 4.18E - 01 | 8.23E - 01 |
| **cKO - M** | 5.98E - 01 | 1.95E - 02 | 5.15E - 01 | 4.23E - 04 |
| **AL** | 5.42E-01 | 9.23E-06 | 4.47E-01 | 7.38E-02 |
| **CR-12h-day** | 5.25E-01 | 2.92E-02 | 1.52E-01 | 1.04E-01 |
| **CR-12h-night** | 8.97E-01 | 4.76E-02 | 7.36E-01 | 8.86E-01 |
| **CR-2h-day** | 8.32E-01 | 4.29E-03 | 3.22E-01 | 2.95E-01 |
| **CR-2h-night** | 7.45E-01 | 1.03E-02 | 9.03E-02 | 7.96E-01 |
| **CR-spread** | 9.41E-01 | 1.19E-02 | 4.72E-01 | 2.79E-01 |

**Table S4: Phase-locking value of mean RE/activity with synodic phase.**

| Data | ΔDays (Date of behavioral data peak − Date of nearest full or new moon) | ΔPhase | Phase-locking value (PLV) | PLV p-value |
|---|---|---|---|---|
| **D1** | 1.09 | 0.22 | 0.76 | 8.00E-04 |
| **D2** | 0.64 | 0.23 | 0.61 | 1.66E-02 |
| **D3** | -0.71 | -0.11 | 0.74 | 5.00E-04 |
| **D4** | -0.85 | -0.18 | 0.74 | 1.00E-04 |
| **D5** | -0.67 | -0.15 | 0.95 | 4.10E-02 |
| **D6** | 1.50 | 0.26 | 0.77 | 6.32E-02 |
| **R1 - F** | -0.25 | -0.12 | 0.66 | 3.00E-04 |
| **R1 - M** | 0.30 | 0.03 | 0.55 | 2.30E-03 |
| **R2 - F** | -0.67 | -0.18 | 0.67 | 2.00E-04 |
| **R2 - M** | -0.22 | -0.06 | 0.77 | 0.00E+00 |
| **Cre - F** | -1.15 | -0.24 | 0.71 | 8.00E-04 |
| **Cre - M** | 1.00 | 0.19 | 0.75 | 0.00E+00 |
| **FxFx - F** | 0.00 | 0.02 | 0.75 | 7.00E-04 |
| **FxFx - M** | -0.67 | -0.20 | 0.78 | 2.00E-04 |
| **Het - F** | -1.00 | -0.22 | 0.69 | 6.17E-02 |
| **Het - M** | -0.82 | -0.23 | 0.75 | 1.20E-03 |
| **cKO - F** | -0.89 | -0.21 | 0.68 | 1.30E-02 |
| **cKO - M** | -1.77 | -0.41 | 0.79 | 4.00E-04 |
| **AL** | -0.15 | -0.02 | 0.81 | 0.00E+00 |
| **CR-12h-day** | 0.09 | 0.01 | 0.77 | 0.00E+00 |
| **CR-12h-night** | -0.13 | -0.04 | 0.74 | 0.00E+00 |
| **CR-2h-day** | -0.77 | -0.19 | 0.77 | 0.00E+00 |
| **CR-2h-night** | -0.42 | -0.11 | 0.75 | 0.00E+00 |
| **CR-spread** | 0.49 | 0.10 | 0.80 | 0.00E+00 |
| **Mean** | **-0.25** | **-0.07** | **0.74** | |
| **SD** | **0.75** | **0.17** | **0.07** | |

**Table S5: Phase-locking value of mean RE/activity peaks with peaks in geomagnetic fluctuations.**

| Data | ΔDays = (Date of behavioral data peak − Date of nearest geomagnetic peak) | Mean phase angle | Phase locking value (PLV) | PLV p-value |
|---|---|---|---|---|
| **D1 (Dresden)** | 3.00 | 0.06 | 0.75 | 1.00E-04 |
| **D2 (Dresden)** | 3.73 | 0.18 | 0.80 | 1.00E-04 |
| **D3 (Dresden)** | 0.07 | 0.01 | 0.71 | 0.00E+00 |
| **D4 (Dresden)** | 0.80 | 0.17 | 0.81 | 0.00E+00 |
| **D5 (Dresden)** | 2.86 | 0.65 | 0.87 | 3.10E-03 |
| **D6 (Dresden)** | 1.86 | 0.47 | 0.84 | 8.80E-03 |
| **R1 - F (Dresden)** | 2.35 | 0.54 | 0.76 | 0.00E+00 |
| **R1 - M (Dresden)** | 1.18 | 0.32 | 0.57 | 3.40E-03 |
| **R2 - F (Dresden)** | -0.44 | -0.07 | 0.84 | 7.00E-04 |
| **R2 - M (Dresden)** | 1.53 | 0.35 | 0.77 | 0.00E+00 |
| **Cre - F (Evanston)** | 0.62 | 0.15 | 0.74 | 6.00E-04 |
| **Cre - M (Evanston)** | 1.44 | 0.35 | 0.75 | 3.00E-03 |
| **FxFx - F (Evanston)** | 0.29 | 0.07 | 0.72 | 7.00E-04 |
| **FxFx - M (Evanston)** | 1.60 | 0.37 | 0.89 | 0.00E+00 |
| **Het - F (Evanston)** | -0.75 | -0.22 | 0.53 | 1.16E-01 |
| **Het - M (Evanston)** | -0.54 | -0.10 | 0.70 | 1.00E-03 |
| **cKO - F (Evanston)** | 2.09 | 0.52 | 0.75 | 7.00E-04 |
| **cKO - M (Evanston)** | 0.29 | 0.06 | 0.76 | 2.00E-04 |
| **AL (Dallas)** | -0.73 | -0.20 | 0.61 | 0.00E+00 |
| **CR-12h-day (Dallas)** | 0.40 | 0.09 | 0.77 | 0.00E+00 |
| **CR-12h-night (Dallas)** | -0.41 | -0.10 | 0.73 | 0.00E+00 |
| **CR-2h-day (Dallas)** | -0.40 | -0.09 | 0.68 | 0.00E+00 |
| **CR-2h-night (Dallas)** | -0.89 | -0.19 | 0.80 | 0.00E+00 |
| **CR-spread (Dallas)** | 0.28 | 0.04 | 0.77 | 0.00E+00 |
| **Mean** | **0.84** | **0.14** | **0.75** | |
| **SD** | **1.31** | **0.25** | **0.08** | |
